# Open-source, high-throughput targeted in-situ transcriptomics for developmental biologists

**DOI:** 10.1101/2023.10.10.561689

**Authors:** Hower Lee, Christoffer Mattsson Langseth, Sergio Marco Salas, Andreas Metousis, Eneritz Rueda Alana, Fernando Garcia-Moreno, Marco Grillo, Mats Nilsson

**Affiliations:** Science for Life Laboratory, Department of Biochemistry and Biophysics, Stockholm University, 171 65, Solna, Sweden; Achucarro Basque Center for Neuroscience, Scientific Park of the University of the Basque Country (UPV/EHU), 48940, Leioa, Spain; Department of Neuroscience, Faculty of Medicine and Odontology, UPV/EHU, Barrio Sarriena s/n, 48940 Leioa, Bizkaia, Spain; IKERBASQUE Foundation, María Díaz de Haro 3, 6th Floor, 48013 Bilbao Spain

**Keywords:** Spatially-resolved transcriptomics, Multi-omics, Multiplexed-FISH

## Abstract

Multiplexed spatial profiling of mRNAs has recently gained traction as a tool to explore the cellular diversity and the architecture of tissues. We propose a sensitive, open-source, simple and flexible method for the generation of in-situ expression maps of hundreds of genes. We exploit direct ligation of padlock probes on mRNAs, coupled with rolling circle amplification and hybridization-based *in situ* combinatorial barcoding, to achieve high detection efficiency, high throughput and large multiplexing. We validate the method across a number of species, and show its use in combination with orthogonal methods such as antibody staining, highlighting its potential value for developmental biology studies. Finally, we provide an end-to-end computational workflow that covers the steps of probe design, image processing, data extraction, cell segmentation, clustering and annotation of cell types. By enabling easier access to highthroughput spatially resolved transcriptomics, we hope to encourage a diversity of applications and the exploration of a wide range of biological questions.

## Introduction

A plethora of methods for multiplexed spatial profiling of mRNAs have recently emerged as tools to explore and visualize cellular diversity in its spatial context (1–6). Some of these methods are sequencing-based and untargeted, hence they can be easily adapted to new model organisms. Other methods are instead imaging-based, and generally targeted, requiring a pre-selection of transcripts whose expression needs to be captured using specific probes. Whether targeted or untargeted, spatial-omics methods often require specialized or proprietary equipment, and their commercial implementations are usually bundled within expensive automated machines and kits, limiting the range of potential uses and applications, and the wider adoption by a more diversified research community.

Targeted *in situ* sequencing (ISS) has been developed as a method for multiplexed mRNA detection (5–7). ISS relies on the ligation of barcoded padlock probes (PLPs) on in-situ synthesized cDNAs, followed by rolling circle amplification (RCA), to generate gene-specific amplicons in situ. These amplicons are large and stable, and can be interrogated by iterative cycles of hybridisation and stripping of fluorescently labeled oligonucleotides, producing very bright signals that can be imaged at low magnification. If using a combinatorial detection scheme, the number of detectable genes scales according to the rule x^N where x is the number of available fluorophores and N is the number of imaging cycles.

A feature of ISS, compared to other multiplexed in-situ methods, is its relatively low detection efficiency. On one hand this is a desirable property: because only a small subset of the total mRNA molecules are captured by padlock probes and massively amplified, the amplicons can be easily visualized using conventional widefield fluorescence microscopes at relatively low magnification without overcrowding the image with signals. This allows the fast imaging and decoding of large tissue sections (high-throughput). On the other hand, because of the very same reason, detection of transcripts expressed at a low level, or from heavily fixed mRNAs, can occasionally be challenging to detect with ISS. Finally, most imaging-based spatial-omics methods produce large and complex image datasets that are often challenging to mine and analyze.

To address these limitations, we first introduced a new detection chemistry with increased capture efficiency, and validated it on a number of use cases. Second, we compiled an end-to-end analysis pipeline, covering the steps of probe design, image analysis, data mining, decoding, cell segmentation and clustering. We also produced a complete manual to guide new users through an entire ISS experiment, and we detail how to adapt our core analysis to different microscopes or input data format.

We hope this work will encourage a larger and diverse research community to experiment with spatially-resolved omics techniques for addressing a broader range of scientific questions.

## Results and discussion

### RNA-ISS recapitulates known mRNA expression patterns with high sensitivity

With the aim of increasing the detection efficiency of ISS, we introduced a novel detection chemistry based on the direct ligation of padlock probes on their target RNA, avoiding the cDNA synthesis step (Fig 1A and Fig 1B).

**Fig. 1.**
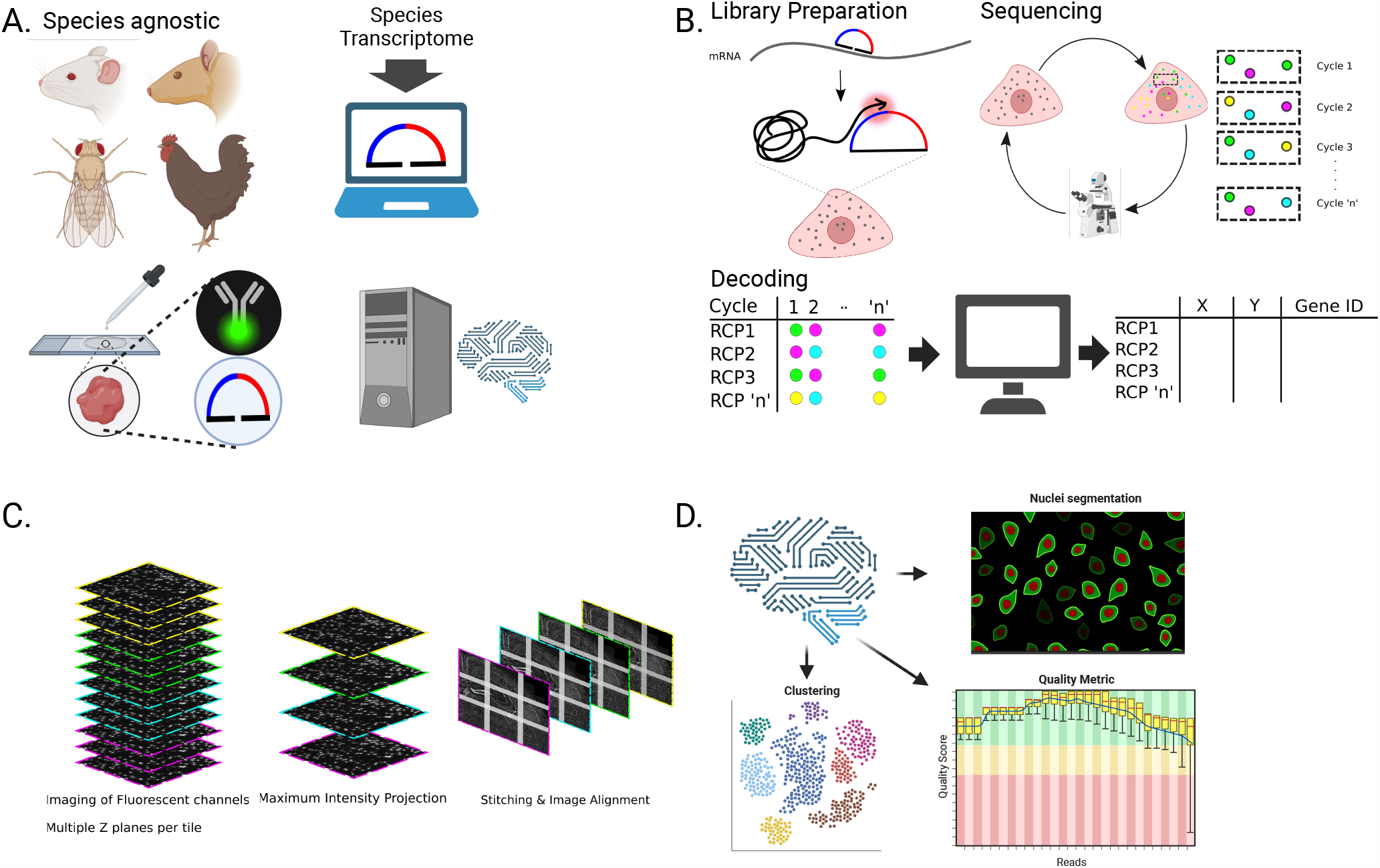
Open-source, high-throughput targeted in-situ transcriptomics: (A) The open-source RNA-ISS assay is species agnostic and is compatible with IHC and other fluorescent labeling, after transcript detection. The workflow includes a custom probe design and packaged processing pipeline. (B) Overview of RNA-ISS. First, gene-specific chimeric padlock probes hybridize to their complementary mRNA target sequence, before they are ligated and amplified by rolling circle amplification. Next, the in situ generated rolling circle products are combinatorially labeled and imaged across multiple cycles and computationally decoded to identify the corresponding genes. (C) Overview of pre-processing pipeline: raw images acquired from the microscope are transformed into a format suitable for decoding: first the images are maximum intensity projected, then simultaneously stitched and registered across imaging cycles. Lastly, the aligned stitched images are sliced again into smaller tiles for computationally efficient decoding. (D) Overview of processing functionalities and features built into the pipeline: Packages for padlock probe design, downstream data analysis such as nuclei segmentation, clustering, quality metrics and probabilistic cell typing functionalities are included to empower researchers with an end to end ISS solution from assay design to computational analyses.

This new chemistry uses chimeric padlock probes in combination with T4RNALigase2, as previously suggested for in vitro applications (8). With these modifications, we achieve an average 2 fold increase in sensitivity (Supp. Table 1) with the same number of probes/transcript compared to the old cDNA-ISS chemistry, implying a set of potential advantages: first, more fine-grained information can be extracted from the same tissue, allowing the resolution of a higher number of transcripts per cell. Second, genes with lower expression levels can be more efficiently detected. Third, informative signal density can likely be extracted with a reduced set of probes, which might be useful for particular applications (detection of short mRNAs, isoform discrimination, etc…). We tested the robustness of the method by applying it, for validation purposes, to several animal model species: *D. melanogaster, M. musculus, G. gallus*, with essentially no modifications to the protocol. Initially we applied the method to the fly ovary, probing for a set of germ-line and somatic mRNAs, and visualizing the amplicons corresponding to each gene with a single fluorescent detection oligo. This allows us to disentangle potential artifacts of the chemistry from issues that might arise from the downstream computational decoding. As expected, *nanos* (Fig 2A) and *vasa* (Fig 2B) are detected exclusively in the nurse cells and in the oocyte (germ-line), *traffic-jam* (Fig 2C) is exclusive to the follicular epithelium (somatic), whereas *slow border cells* (Fig 2E) is exclusive to a subgroup of somatic cells called border cells. Our method also accurately resolves the subcellular localization of the *gurken* transcript, showing a dense signal accumulation surrounding the dorso-anterior side of the oocyte nucleus, where it is expected at this oocyte maturation stage and indicating that our method accurately preserves the subcellular localization of mRNAs (Fig 2D).

**Fig. 2.**
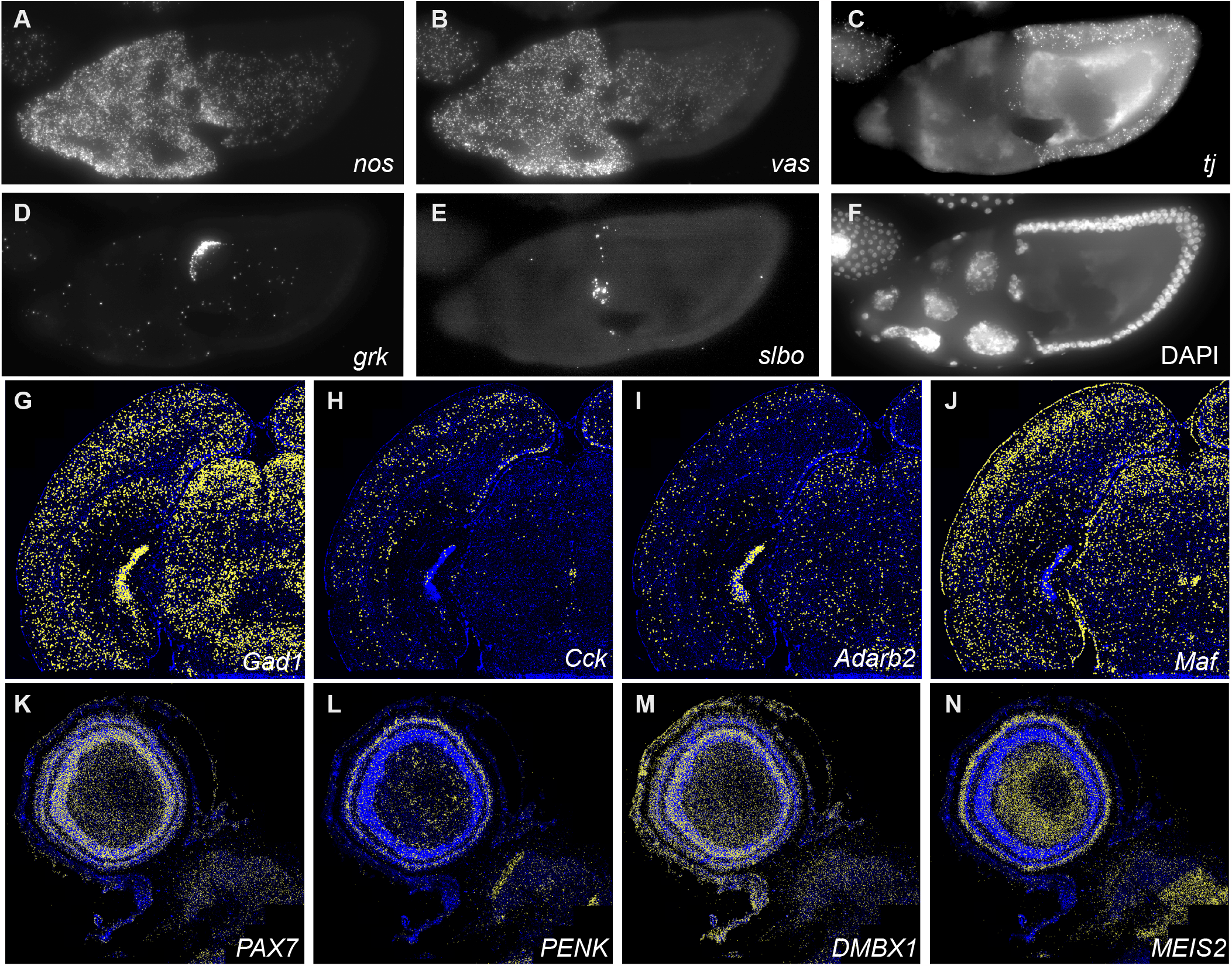
RNA-ISS recapitulates known mRNA expression patterns: (A-F) Fluorescent signal detected by direct binding of detection oligos to the rolling circle products for several transcripts on the *Drosophila* ovary. Germline specific genes (*vasa* and *nanos*) are detected in the nurse cells and the oocyte (panels A and B respectively), and are excluded in the follicular epithelium, which is generally labeled by the expression of *traffic-jam* (panel C). (D) The antero-dorsal subcellular localization of *gurken* in the oocyte is accurately reproduced. (E) the somatic sub-population of border cells expresses *slobo*, a known marker. (F) DAPI counterstaining (G-J) Expression of a set of markers in a coronal section of the mouse brain. (G) Gad1, a GABAergic neuron marker, is expressed by inhibitory neurons in the cortex, hippocampus, thalamus and superior colliculus, as well as in the midbrain tegmentum; (H-I) *Cck* and *Adarb2*, both interneuronal marker, label interneurons in the cortex and hippocampus; *Adarb2* also shows enriched expression in the telencephalic choroid plexus, and scattered expression in the midbrain; (J) *Maf*, an inhibitory neuronal marker is expressed in the cortex, hippocampus, superior colliculus and thalamus, and highly enriched expression appears in the periaqueductal grey matter - all recapitulating known spatial expression pattern, as validated with ISH data from the Allen Brain Atlas and mousebrain.org. (K-N): Expression of a set of markers in the optic tectum (OT) of the E15 chicken brain. Different markers specifically highlight cells belonging to specific layers of the OT. (K) *PAX7*, a transcription factor expressed in pretectum and mesencephalon appears enriched in all tectal layers; (L) neuropeptide-encoding gene *PENK* is expressed in superficial tectal layers and in geniculate pretectal nucleus; (M) *DMBX1*, a transcription factor typical of the diencephalon and mesencephalon, shows ubiquitous expression in all layers and meninges. (N) transcription factor *MEIS2* shows highest expression in the stratum griseum centrale near the ventricle and in superficial strata. All these expression patterns recapitulate expression patterns found in the literature and Geisha Arizona database.

We then designed two further probe sets, respectively against genes expressed in mouse coronal sections (15 genes) and sections of the chicken optic tectum (35 genes), proceeded to apply our detection chemistry, and computationally decoded their expression. For both species, the decoded expression patterns are localized, specific, and consistent with previous available knowledge (Figure 2 G-J for mouse and Figure 2 K-N for chicken).

### RNA-ISS is compatible with standard labeling techniques

We reasoned that the ability of combining routine assays with ISS would be of crucial interest to many developmental biologists, so we tested the compatibility of our detection chemistry with a few labeling techniques commonly used in the field. To test whether ISS could be combined with immunostaining, after the last cycle of ISS detection on mouse brains we stripped all the detection probes using formamide, and proceeded to stain against the GFAP protein, successfully recapitulating the expected specific pattern (Figure 3 H-O). This is of potential interest for different reasons: 1) it allows researchers to correlate ISS data with known landmarks on the tissues, marked by the specific expression of a given protein; 2) in samples for which multiplexed antibody panels are available (ie. human), high-plex multi-modal interrogation of a tissue might be possible; 3) simultaneous analysis of mRNA and proteins encoded by the same set of genes potentially allows the study of mRNA translation dynamics, and finally 4) staining for membrane proteins might allow a rigorous transcript segmentation. We then tested the possibility of combining ISS with EdU-labeling, for simultaneous spatial analysis of gene expression and birthdating: we injected EdU into E4 developing chicken, sacrificed the embryo at E15, probed a gene set that would allow us to classify cell types in the diencephalon, and successfully revealed the EdU labeling by click chemistry at the end of the process. The results from these experiments, shown in the accompanying paper/preprint (9), indicate that ISS and EdU-based birthdating can be easily combined on the same tissue sample, allowing to interrogate both gene expression and timing of origin of selected cell types within a tissue. These two examples of how ISS can be integrated with existing techniques showcase only a relatively small range of potential applications. Other ideas that come immediately to mind include the combined use of ISS with genetically-encoded photoconvertible sensors (ie, CaMPARI, (10) or with clonal tracing tools such as CRE-LOX (11) or FLP-FRT (12).

**Fig. 3.**
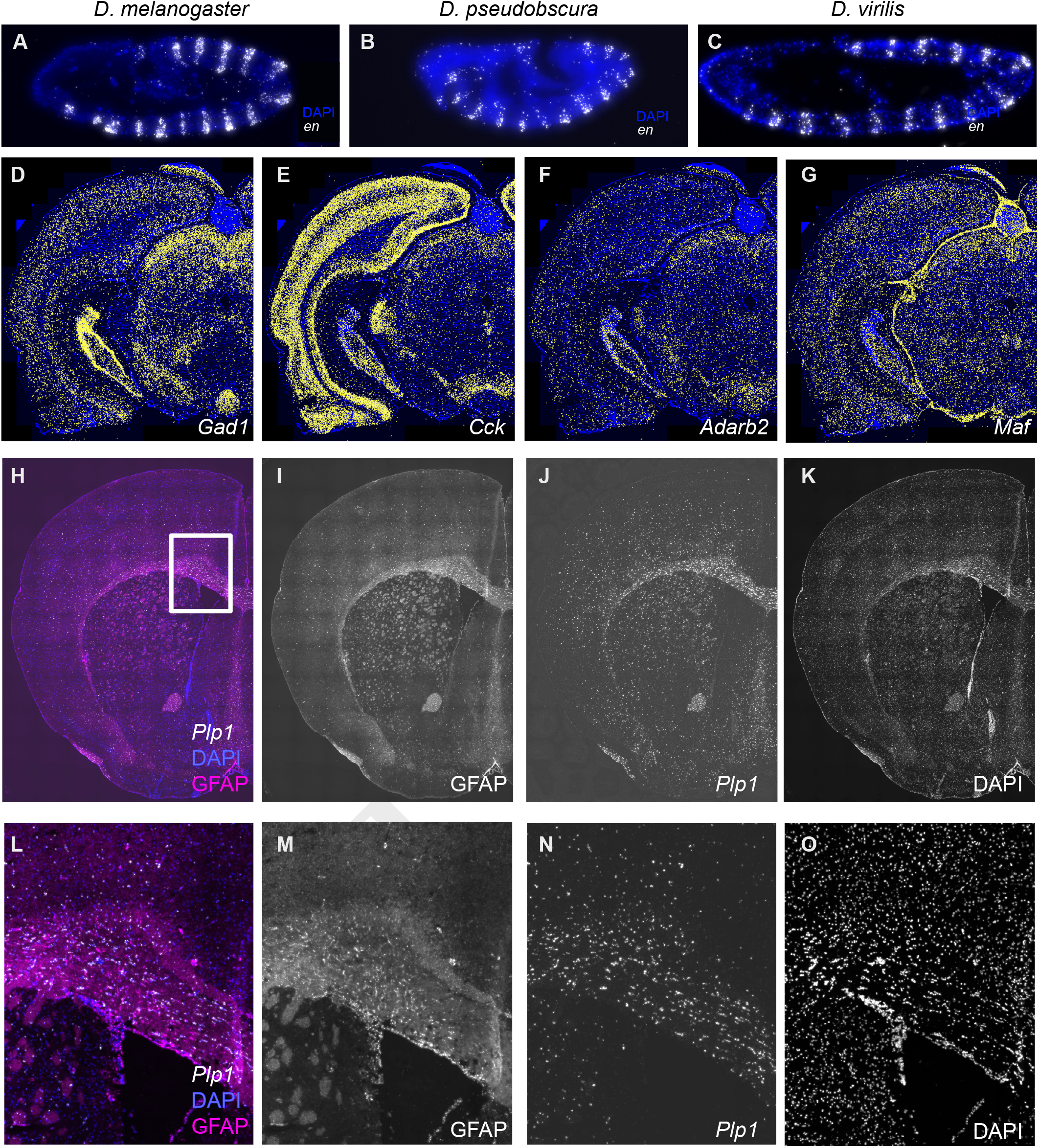
RNA-ISS enables cheap comparative analysis in related species and is compatible with antibody labeling: (A-C) A set of probes targeting conserved regions are able to detect the mRNA of *engrailed* across multiple related species, *D. melanogaster* (A), *D. pseudobscura* (B), and *D. virilis* (C), with comparable efficiency. The species cover an evolutionary divergence time of approximately 40 million years.(D-G) The same cross-reactive design was used to produce mouse-rat cross reactive probes. The same genes of Fig. 2 (D-G) are probed on rat brains, using the probes designed against mouse, showing similar expression patterns. (H-O) RNA-iss can be combined with posterior (or simultaneous) immunohistochemistry. (H) After performing ISS, the detection probes were stripped and the tissue stained using an antibody against the GFAP protein (I). The expression of the *Plp1* mRNA was also re-labeled with a fluorescent detection oligo (J), to provide a contrast reference. (K) DAPI counterstaining. (L) magnified inset spanning the area highlighted with a white rectangle in panel H, respectively for panel H. (M-O) show the isolated channels (respectively GFAP, *Plp1* and DAPI for the inset image).

### Cross-reactive design of probes allows low-cost comparative analysis of gene expression

A peculiar feature of the RNA-based ISS chemistry is its mismatch-tolerance. While the direct probing of RNA produces an overall increase in detection efficiency, it also generates a small specificity cost, because the enzyme of choice is, to some extent, mismatch tolerant (8). As a result, it is sometimes impossible to design padlock probes to discriminate between the mRNAs of closely related genes (ie: recent gene duplications), because their sequence has not diverged enough to fall above the ligase specificity threshold. We seeked to turn this limitation into an advantage, by rationally designing cross-reactive padlock probes that are promiscuous enough to recognise the same gene’s mRNA across multiple related species, thus enabling comparative analysis of gene expression with a reduced experimental cost. We tested this approach by designing a set of PLPs against the engrailed gene of *D. melanogaster*, explicitly selecting targets with a high sequence conservation among the “12 Drosophila species” set. Besides *D*.*melanogaster*, this probe set shows specific and sensitive activity on *D. pseudobscura* and *D. virilis* embryos, showing that this design can be exploited to experimentally cover potentially large evolutionary periods: the last common ancestor of *D. melanogaster* and *D. virilis* is estimated to be 40 MYA, a time interval corresponding roughly to the split between old and new world monkeys. We tested the same design logic on a set of genes expressed in mouse and rat midbrain coronal sections, using a set of padlock probes explicitly designed to work in both species (<20 MY apart), showing that our strategy applies also for this case. Finally, we predicted in silico the cross-reactivity on *Macaca fascicularis* of a large panel of 1776 probes previously designed against 363 human genes. These probes were not explicitly designed to be cross-reactive, however we found that roughly 74% of them is predicted to recognise specifically individual genes in *Macaca*, 12% is predicted not to bind to any transcript, while 14% is predicted to recognise off-target mRNAs. The rational design of cross reactive probes is particularly interesting for labs focused in comparative work, allowing the spatial analysis of large numbers of genes across multiple related species.

### A toolbox for design and data analysis of ISS experiments

To enable naive users to easily design reagents for RNA-based ISS, process the imaging data and analyze the experimental results, we compiled a set of Jupyter notebooks and installable packages that can be easily followed, as well as an accompanying step-by-step manual to guide the user through the whole process of image preprocessing and analysis (supplementary manual). We give here a brief overview of the main functionalities of each notebook, but we refer to the supplementary manual for more detailed information and guidance.

Probe design: given a set of gene IDs and a transcriptome of reference, the software helps the user extract a number of suitable target regions from each mRNA sequence, and checks these regions for off-targets in the transcriptome. The script also associates each gene to a unique barcode and assembles the final padlock probe sequences. Preprocessing: After the wet lab and imaging parts are completed (see Materials and Methods), the images are fed into the preprocessing module. This module transforms the raw images into a format suitable for decoding, executing the following steps: 1) the images are maximum Z-projected. 2) the resulting 2D projected images (“tiles”) are simultaneously stitched and registered across imaging cycles, using the ASHLAR software (13). 3) the aligned stitched images are sliced again into smaller tiles, to allow a faster and computationally efficient decoding. Decoding: The preprocessed images are fed into the decoding module, where they get converted into SpaceTX format (14), and piped into the Starfish Python library for decoding of image-based spatial transcriptomics datasets (https://github.com/spacetx/starfish). The output of the decoding is a csv table, where each row represents a mRNA detection event, and different properties of each spot are extracted (x and y position, corresponding gene, quality metrics, etc…). A set of custom functions allows the users to interactively generate plots and figures for individual genes, as well as to inspect some of the performance metrics of the decoding process. Postprocessing: The decoded transcript table can be fed to the postprocessing module. Here, several functions enable the user to segment cells using some published algorithms, Cellpose (15)] and Stardist (16), assign transcripts to cells and finally create Annotated Data objects (AnnData), for more advanced analysis using the scverse ecosystem of tools (17). New users wishing to repurpose an available microscope to perform ISS would essentially need to interface the output images from such microscope with the provided preprocessing module. Once the preprocessing step is correctly completed, the other modules are expected to work with minimal modification. We detail in the manual the requirements of the input files for the preprocessing module.

Summing up, we provide an open-source method for highsensitivity in situ sequencing that is readily applicable to a wide range of tissues types and animal model organisms, both vertebrates and invertebrates. We believe the method could be easily transferred to other species (including non-animals) with minimal modification. The wet lab workflow is easily implemented on top of standard procedures such as immunostaining and EdU labeling, making it particularly valuable for developmental biologists. We also provide a simplified bioinformatic pipeline for data mining, decoding, and advanced analysis of ISS datasets, from raw microscope images to cell clusters, as well as a complete guide to the entire analysis workflow. We hope this work encourages the adoption of imaging based spatial-omics by a large and diverse community of scientists in the developmental biology field.

## Material and methods

### Probe design, synthesis, pooling and phosphorylation

The notebook for probe design provided at (https://github.com/Moldia/Lee_2023/) performs the following operations: given a list of gene IDs and a transcriptome of reference, the mRNAs corresponding to all the described isoforms of each gene are extracted and aligned using CLUSTALW2 (18). The common regions are then sliced in all the possible 30-mers that compose them (nt 1-30, 2-31, etc…), and the resulting 30mers are filtered according to increasingly restrictive criteria to find suitable targets for the binding of the padlock probes: first, a GC content filter is applied, to retain only 30mers within a given range. Among these, only the 30mers with a C or G in position 16 are kept: this is because T4RNAl2 has a slight positive efficiency bias towards terminal 3’ G or C. This constraint can be easily removed to facilitate probe design on difficult targets. A random subset of non-overlapping suitable 30mers spanning the transcript is then “specificity-checked”: each target is searched against the transcriptome, using the pattern-matching capability of Cutadapt (Martin, 2011) and allowing for up to 6 mismatches, a conservative threshold under which we expect at least some degree of cross-reactivity. 30mers showing non-specific hits (ie: hits that do not belong to the query gene) are discarded: because T4RNAl2 is slightly promiscuous, performing this step ensures that the PLPs will have no off-targets. Of all the 30mers passing the specificity check, a subset (normally 5) is chosen, and a final step creates a specific padlock probe sequence in which unique barcodes are associated with unique genes. To design explicitly cross-reactive probes for multiple fly species, we performed the 30mer step extraction for all the *engrailed* orthologues across the 12 *Drosophila* species. We then kept for the downstream steps only the 30mers that had a difference of less than 6 nucleotides from the *D. melanogaster sequence*. Finally, we filtered out all the 30mers that could produce secondary hits in the transcriptome, using our specificity check based on Cutadapt. Similarly, for mouse and rat cross reactivity, we ran the 30mer search in both species, and retained only the 30mers found in both species, with a tolerance of 6 mismatches. Of these, hits with potential off targets were excluded in the posterior filtering steps.

To assess the possible cross-reactivity of a human probe panel on *Macaca*, we run a specificity check for the human probes with a 6 nucleotide mismatch tolerance. We then classified the probes in 3 categories, according to the results: a) probes that recognize the mRNA of a single specific gene (“cross reactive and specific”); b) probes that recognize the mRNAs of more than one gene (“cross reactive but non-specific”); c) probes that do not recognize any mRNA (“non-cross reactive”).

The probes were ordered as DNA Ultramers with a 3’ terminal RNA base from Integrated DNA Technologies (IDT), with a synthesis scale of 4nmol, resuspended in IDTE buffer at a 200 uM concentration. Probes can either be ordered prephosphorylated (with some added cost) or phosphorylated in house. When working with large pools, we used the latter approach. We pooled all the padlock probes for a given experiment in a single tube, and phosphorylated according to the following protocol: 20 nmol of the pooled probes were phosphorylated in 1X PNK buffer (B0201S, New England Biolabs), 1 mM ATP (P0756S, New England Biolabs) and 20 units of T4 Polynucleotide kinase (M0201L, New England Biolabs), in a total volume of 50 uL, for 2 hours at 37 degrees C. PNK was then heat-inactivated at 65C for 5 minutes and the pooled phosphorylated probes were stored at −20C until used.

### Tissue fixation and pre-processing

Mouse and chicken brains were dissected and immediately immersed in OCT embedding medium (361603E, VWR) for a washing step, and then transferred to an embedding mold containing OCT for embedding and freezing. The embedding mold was transferred to a dry-ice box and frozen for 10 minutes (we found that this freezing method produces better tissue integrity than liquid nitrogen). Samples were stored at −80 until sectioning. Embedded tissue blocks were sectioned on a cryostat in slices 10 to 20 micrometers thick and attached to Superfrost Plus microscope slides (631-0108, VWR). The sections can be stored at −80 indefinitely. The first day of the library prep protocol, we thawed the samples for 5 minutes allowing them to reach room temperature, washed the slides in PBS and immersed them in PFA 3% for 5 minutes. This is the starting point of the library prep protocol referred below when working with fresh frozen material.

For *Drosophila tissues*, we collected overnight egglays on apple juice agar plates, dechorionated the embryos with bleach, and fixed them in a 1:1 mix of 4% Formaldehyde in PBS and heptane. We then removed the heptane, devitellinized the embryos using methanol, and stored them until the day of embedding. Similarly, ovaries were dissected on ice-cold PBS and immediately fixed in 4% formaldehyde for 20 minutes, then washed with PBS 3 times, dehydrated progressively in methanol:PBS (1:3, 2:2, 3:1, 20 minutes each), washed twice in methanol and stored in methanol at −20 C until embedding. For embedding fly tissues, we rehydrated them progressively in methanol:PBS (3:1, 2:2,1:3, 20 minutes each) and washed twice in PBS. We then cryoprotected the tissues overnight with 30% sucrose in PBS and finally transferred the tissues to OCT-containing embedding molds, and froze the samples on dry ice for 10 minutes. Samples were stored at −80 until sectioning. Embedded tissue blocks were sectioned on a cryostat in slices 10 to 20 micrometers thick and attached to Superfrost Plus microscope slides. The sections can be stored at −80 indefinitely. The first day of the library prep protocol, we thawed the samples for 5 minutes allowing them to reach room temperature, and washed them with PBS.This is the starting point of the library prep protocol referred below when working with PFA-fixed material.

Rat brain slices were purchased fresh-frozen (Zyagen), kept at −80 until use and processed the same way as chicken and mouse sections.

### RNA-ISS Library prep

Fixed slides were washed 3 times with PBS (room temperature) and permeabilized with a 0.1 N HCl incubation for 5 minutes, followed by 2 washes in PBS. The samples were progressively dehydrated with a 70% ethanol bath for 2 minutes, followed by a 100% ethanol bath for 2 minutes, then air dried. We attached secure-seal chambers to cover the samples, and filled the chamber with PBS-Tween 0.5%, followed by a PBS wash. A probe solution was prepared according to the following recipe: 2x SSC, 10% Formamide and 10 nm of each padlock probe, and incubated on the samples overnight at 37 C. The next day we washed the unhybridized excess probes by 2 washes of 10% formamide in 2x SSC, followed by 2 washes in 2x SSC. After removing the last SSC wash, a ligation mix was prepared as described in the “Probe ligation” step in the attached protocol and incubated on the samples for 2 hours at 37 degrees Celsius. After ligation, the samples were washed twice with PBS and an amplification mix was prepared as in the “Amplification of the padlock probes” step in the attached protocol. The amplification reaction was carried out overnight at 30 C. The next day, the samples were washed 3 times with PBS and L-probes (or bridge probes) were incubated for 30 minutes as described in the step “hybridization of L-probes” in the attached protocol. Excess probes were washed out with 2 washes in 2x SSC and detection oligos and DAPI were incubated for 30 minutes as indicated in the protocol. Excess detection oligos were washed out with 2 washes in 2x SSC. If necessary, TrueBlack was applied to quench background fluorescence, according to the manufacturer’s instructions. Samples were mounted in SlowFade gold, and cyclical imaging was performed. After each cycle of imaging, Lprobes and detection oligos were stripped with two washes of 3 minutes in 100% formamide, followed by 5 washes in 2x SSC. Hybridisation of L-probes and detection oligos for the following detection cycle was performed as above. A step by step protocol walk-trough is available at: https://www.protocols.io/edit/home-made-direct-rna-detection-cqguvtww

### Image acquisition

Imaging was performed using a standard epifluorescence microscope (Leica DMI6000) connected to an external LED source (Lumencor® SPECTRA X light engine). Light engine was set up with filter paddles (395/25, 438/29, 470/24, 555/28, 635/22, 730/40). Images were obtained with a sCMOS camera (2048 × 2048, 16-bit, Leica DFC90000GTC-VSC10726), automatic multislide stage, and Leica Apochromat objectives 20x (20X, HC PL APO 20x/0.80 DRY, 11506529), 40× (HC PL APO 40x/1.10 WATER, 11506342). The microscope was equipped with filter cubes for 6 dye separation (AF750, Cy5, Cy3, AF488, Atto425 and DAPI) and an external filter wheel (DFT51011).

Each region of interest (ROI) was marked and saved in the Leica LASX software, for repeated imaging. Each ROI was automatically subdivided into tiles, and for each tile a z-stack with an interval of 0.5 micron was acquired in all the channels. The tiles are defined to have a 10% overlap at the edges. The images were saved as thousands of individual tiff files with associated metadata.

### Data processing

The raw images and the associated meta-data from the microscopes were fed into the preprocessing module of our analysis pipeline. The module transforms the images into a format suitable for decoding, executing the following steps: first the images are maximum Z-projected. The resulting 2D projected images (“tiles”) are simultaneously stitched and registered across imaging cycles, using the ASHLAR (13) software. ASHLAR captures the metadata and places the tiles correctly in the XY space before starting the alignment step. During the process, the 10% overlap is also removed to produce stitched images. Finally, the aligned stitched images are sliced again into smaller tiles, to allow a faster and computationally efficient decoding. The resliced aligned images then taken over by the decoding module, which converts them into the SpaceTX format (14), and pipes them into the Starfish Python library for decoding of image-based spatial transcriptomics datasets (https://github.com/spacetx/starfish). Within the decoding modules, the images are normalized across channels and imaging cycles, a spot detection step is performed and for each detected spot its intensity across all channels is extracted. For each spot, the prominent channel for each cycle is extracted and a spot identity is annotated in color space. Each spot is now represented by a sequence of colors across cycles. The color sequence of each spot is matched to a decoding table that associates a color sequence with a specific gene. The output of this decoding is a csv table, in which each row represents a detection spot with different properties (x and y position, gene identity, quality metrics). The quality metrics for each spot is computed as follows: for each spot, the normalized fluorescence intensities across all channels are extracted. The prominent channel is considered the “true signal”, while all the others are considered “background”. The score is described by the formula “true signal” / (“true signal” + “background”), and it has a theoretical maximum of 1 (perfect decoding) and a theoretical minimum value of 0.25 when decoding in 4 colors, which corresponds to a random assignment (ie. all the channels have the same fluorescence intensity for that spot). The quality score for each spot is computed per cycle, allowing to calculate 2 parameters: 1) the average quality across all cycles and, 2) the minimum quality across cycles. We found that filtering according to a minimum quality produces more reliable data, and we normally use a filter value of 0.5. For the chicken optic tectum sections, we additionally performed image deconvolution using *flowdec* (19) on the raw images before proceeding to the preprocessing steps. We find this step to drastically increase the number of detected spots in dense datasets. Other modules in the repository allow users to perform additional operations on the images, and are documented in the supplementary manual.

### Antibody staining

After RNA-ISS detection, tissue was washed 3 times in 100% formamide for 2 minutes, to remove hybridized detection probes, and washed 5x in PBS. Then the tissue was blocked with PBTA (PBS, 5% normal donkey serum (Jackson ImmunoResearch), 0.5% Triton-X 100) for one hour. Then sections were incubated with primary antibodies, GFAP (Dako, Z0334) overnight at +4 °C. Sections were then washed three times with PBS and incubated with secondary antibodies (Alexa Fluor 488 anti-rabbit) for 2 h at room temperature and counterstained with DAPI.

## Supporting information

Supplementary Manual

## Code availability

All the code used in this paper is available at https://github.com/Moldia/Lee_2023.

## Author contributions

HL performed most of the chemistry optimization and the mouse-rats experiments. CML and SMS wrote most of the analysis code. AM designed the scripts for cross-reactive probes. ERA performed the chicken experiments, supervised by FGM who also contributed the interpretation of expression data. MG performed early tests, contributed to the code, conceptualised the study and wrote the paper. MN directed and supervised the work, secured funding, and provided critical feedback on the work and the text.

## ACKNOWLEDGEMENTS

We’d like to thank the members of the Nilsson lab, of the In-situ sequencing platform at Scilifelab, and of the imaging facility at EMBL Monterotondo (particularly Alvaro Crevenna and Alejandro Linares), for useful discussion and extensive testing of the chemistry, as well as for their constructive feedback on the software. The work in MN’s group is supported by funds from by Chan Zuckerberg Initiative, an advised fund of Silicon Valley Community Foundation; Erling-Persson Family Foundation (A human developmental cell atlas); Knut and Alice Wallenberg Foundation (KAW 2018.0172); Swedish Research Council (2019-01238) and Swedish Cancer Society (Cancerfonden; CAN 2021/1726). ERA holds a predoctoral fellowship from the Basque Government. During the duration of this research, FGM holds and held an Ikerbasque Research Fellowship, Spanish Ministry MICNN PGC2018-096173-A-I00 and PID2021-125156NB-I00 grants, Basque Government PIBA 2020_1_0057 and PIBA_2022_1_0027 grants and EASI-GENOMICS 3^rd^ TNA call PID14596 grant.

---

**Table 1.** Efficiency comparison of RNA-ISS vs cDNA.

|  | Signal dots per nucleus |  |  |
| --- | --- | --- | --- |
|  | Plp1 | Rorb | Foxj1 |
| cDNA thalamus | 1.756711 | 0.049269 | 0.03262 |
| cDNA ventricle | 2.480241 | 0.067649 | 0.168118 |
| dRNA thalamus | 2.591968 | 0.211187 | 0.053783 |
| dRNA ventricle | 3.709754 | 0.151536 | 0.268647 |

## Bibliography

1. Lars E Borm, Alejandro Mossi Albiach, Camiel C A Mannens, Jokubas Janusauskas, Ceren Özgün, David Fernández-García, Rebecca Hodge, Francisca Castillo, Charlotte R H Hedin, Eduardo J Villablanca, Per Uhlén, Ed S Lein, Simone Codeluppi, and Sten Linnarsson. Scalable in situ single-cell profiling by electrophoretic capture of mRNA using EEL FISH. Nat. Biotechnol., 41(2):222–231, February 2023.

2. Kok Hao Chen, Alistair N Boettiger, Jeffrey R Moffitt, Siyuan Wang, and Xiaowei Zhuang. RNA imaging. spatially resolved, highly multiplexed RNA profiling in single cells. Science, 348(6233):aaa6090, April 2015.

3. Simone Codeluppi, Lars E Borm, Amit Zeisel, Gioele La Manno, Josina A van Lunteren, Camilla I Svensson, and Sten Linnarsson. Spatial organization of the somatosensory cortex revealed by osmFISH. Nat. Methods, 15(11):932–935, November 2018.

4. A M Femino, F S Fay, K Fogarty, and R H Singer. Visualization of single RNA transcripts in situ. Science, 280(5363):585–590, April 1998.

5. Daniel Gyllborg, Christoffer Mattsson Langseth, Xiaoyan Qian, Eunkyoung Choi, Ser-gio Marco Salas, Markus M Hilscher, Ed S Lein, and Mats Nilsson. Hybridization-based in situ sequencing (HybISS) for spatially resolved transcriptomics in human and mouse brain tissue. Nucleic Acids Res., 48(19):e112, November 2020.

6. Hower Lee, Sergio Marco Salas, Daniel Gyllborg, and Mats Nilsson. Direct RNA targeted in situ sequencing for transcriptomic profiling in tissue. Sci. Rep., 12(1):7976, May 2022.

7. Rongqin Ke, Marco Mignardi, Alexandra Pacureanu, Jessica Svedlund, Johan Botling, Carolina Wählby, and Mats Nilsson. In situ sequencing for RNA analysis in preserved tissue and cells. Nat. Methods, 10(9):857–860, July 2013.

8. Tomasz Krzywkowski, Malte Kühnemund, and Mats Nilsson. Chimeric padlock and ilock probes for increased efficiency of targeted RNA detection. RNA, 25(1):82–89, 2019.

9. Eneritz Rueda-Alaña, Marco Grillo, Enrique Vazquez, Sergio Marco Salas, Rodrigo Senovilla-Ganzo, Laura Escobar, Ana María Aransay, Ana Dopazo, Juan Manuel Encinas, Mats Nilsson, and Fernando García-Moreno. BirthSeq, a new method to isolate and analyze dated cells from any tissue in vertebrates.

10. Benjamin F Fosque, Yi Sun, Hod Dana, Chao-Tsung Yang, Tomoko Ohyama, Michael R Tadross, Ronak Patel, Marta Zlatic, Douglas S Kim, Misha B Ahrens, Vivek Jayaraman, Loren L Looger, and Eric R Schreiter. Neural circuits. labeling of active neural circuits in vivo with designed calcium integrators. Science, 347(6223):755–760, February 2015.

11. B Sauer and N Henderson. Site-specific DNA recombination in mammalian cells by the cre recombinase of bacteriophage P1. Proc. Natl. Acad. Sci. U. S. A., 85(14):5166–5170, July 1988.

12. K G Golic and S Lindquist. The FLP recombinase of yeast catalyzes site-specific recombination in the drosophila genome. Cell, 59(3):499–509, November 1989.

13. Jeremy L Muhlich, Yu-An Chen, Clarence Yapp, Douglas Russell, Sandro Santagata, and Peter K Sorger. Stitching and registering highly multiplexed whole-slide images of tissues and tumors using ASHLAR. Bioinformatics, 38(19):4613–4621, September 2022.

14. Shannon Axelrod, Matthew Cai, Ambrose Carr, Jeremy Freeman, Deep Ganguli, Justin Kiggins, Brian Long, Tony Tung, and Kevin Yamauchi. starfish: scalable pipelines for image-based transcriptomics. J. Open Source Softw., 6(61):2440, May 2021.

15. Carsen Stringer, Tim Wang, Michalis Michaelos, and Marius Pachitariu. Cellpose: a generalist algorithm for cellular segmentation. Nat. Methods, 18(1):100–106, January 2021.

16. Uwe Schmidt, Martin Weigert, Coleman Broaddus, and Gene Myers. Cell detection with Star-Convex polygons. In Medical Image Computing and Computer Assisted Intervention – MICCAI 2018, pages 265–273. Springer International Publishing, 2018.

17. Isaac Virshup, Danila Bredikhin, Lukas Heumos, Giovanni Palla, Gregor Sturm, Adam Gayoso, Ilia Kats, Mikaela Koutrouli, Scverse Community, Bonnie Berger, Dana Pe’er, Aviv Regev, Sarah A Teichmann, Francesca Finotello, F Alexander Wolf, Nir Yosef, Oliver Stegle, and Fabian J Theis. The scverse project provides a computational ecosystem for single-cell omics data analysis. Nat. Biotechnol., 41(5):604–606, May 2023.

18. M A Larkin, G Blackshields, N P Brown, R Chenna, P A McGettigan, H McWilliam, F Valentin, I M Wallace, A Wilm, R Lopez, J D Thompson, T J Gibson, and D G Higgins. Clustal W and clustal X version 2.0. Bioinformatics, 23(21):2947–2948, November 2007.

19. Eric Czech, Bulent Arman Aksoy, Pinar Aksoy, and Jeff Hammerbacher. Cytokit: a single-cell analysis toolkit for high dimensional fluorescent microscopy imaging. BMC Bioinformatics, 20(1):448, September 2019.

