## Supplementary Manual for "Open-source, high-throughput targeted in-situ transcriptomics for developmental biologists"

### Targeted high sensitivity ISS Manual


**Before you start** 3
- How ISS works 3
- Glossary, useful definitions, abbreviations and clarifications: 4
- Imaging and microscope requirements 5
- How ISS analysis works 6
- File naming and formats conventions 7
- Computer requirements 7

**Basic introduction to installing python packages** 8

**The jupyter project** 9
- jupyter lab vs jupyter notebook 9

**Installation of preprocessing, decoding and postprocessing toolboxes** 9

**Run your first experiment:** 10
- Design probes for direct RNA detection. 10
  - Installation of the probe design software tools: 10

**ISS protocol and imaging** 11

**Preprocessing** 12
- Preprocess images from the microscope. 12
  - Objective of the preprocessing 12
  - A note on intermediate files 12
- Running preprocessing 16
  - preprocessing_main_leica 16
    - Explanation of the variables 17
  - Running each of the steps separately 18

**Decoding** 19
- Objective of the decoding 19
- Format images into SpaceTx format 19
  - Explanation of the variables 20
- Extract and decode ISS data from the images. 21
  - What do we do when we run process_experiment? 22
  - Parameters to modify when decoding (process_experiment) 23
  - Combining all decoded tiles into a single file with the decoded spots 24
  - Understand the results and quality metrics. 25
    - How is the quality of the signals calculated? 25
      - What exactly is a "base" in ISS? 25
    - Plotting your quality metrics 25

Filtering the results 30
Plot and visualize the decoded data. 30

**Postprocessing 31**
Cell segmentation 31
Nuclei based cell segmentation 31
cellpose (GitHub - MouseLand/cellpose: a generalist algorithm for cellular segmentation with human-in-the-loop capabilities) 31
Explanation of the variables 32
stardist (StarDist - Object Detection with Star-convex Shapes) 32
Explanation of the variables 33
Pixel based classification 33
Spot based segmentation 33
What can we do downstream of the cell segmentation? 34
Probabilistic cell typing by in situ sequencing (pciSeq) 34
tangram 34
Annotated data objects 34
Create annotated data objects 34
Explanation of the variables 35
Cluster your cells 36
Purpose of clustering 36
When should you do it? 36
Procedure 36
Formatting the data 36
Filtering and normalizing your cells 40
Principal component analysis, KNN and dimensional reduction. 41
Additional functionality in ISS_postprocessing 42
Basic plotting function using sc.pl.spatial 42
Advanced features: 43
Image deconvolution options (requires GPU) 43
Tensorflow-based image deconvolution using Flowdec. 43
Installation and requirements: 43
Data formatting and interfacing with the main pipeline 44
Creating image stacks from Leica-exported tiffs, all cycles (Leica/Nilsson naming only) 44
Creating image stacks from CZI files, all cycles 45
Creating image stacks from Leica-exported tiffs, single cycle (Leica/Nilsson naming only) 45
Creating image stacks from CZI files, single cycle 45
Image deconvolution using flowdec 46
Define PSFs for your microscopes and channels 46
Running flowdec to deconvolve a full experiment 46

ML-accelerated image restoration using 2D CARE: 48
Background 48
Installation and requirements 48
Data formatting and interfacing with the main pipeline 49
Apply a pre-trained model 49
Explanation of the variables 49
Train your own model 50
Automatic processing of CZI files. 50
**References** **51**

### Before you start

#### How ISS works

ISS (In-situ sequencing; (Ke et al. 2013) is a method for highly multiplexed in-situ target transcriptomics analysis developed in Mats Nilsson's lab (http://www.nilssonlab.org).
The method relies on the in situ hybridization and ligation, on endogenous mRNAs, of padlock probes, which are then amplified on the tissue, generating amplicons that can be interrogated ("sequenced") in-situ.

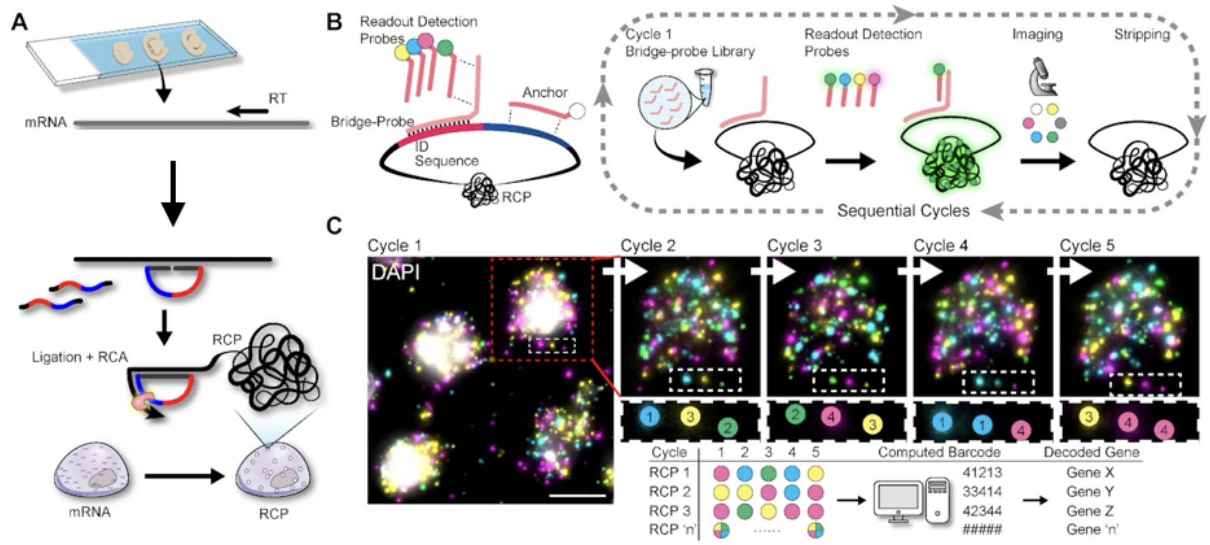


*Figure 1. General scheme of the experimental workflow for ISS. A. Padlock probes are hybridised, ligated and amplified on a tissue. B. Amplicons are detected using a combinatorial labeling strategy across multiple cycles of detection, imaging and stripping. C. Images are computationally processed and the amplicon identity is decoded. Figure taken from (Gyllborg et al. 2020)*

Padlock probes, and hence the amplicons they produce (rolling circle products, RCPs), are typically barcoded according to the mRNA they recognize. These barcodes can be read in a highly multiplexed fashion by iterative combinatorial labeling using fluorescent probes. Using this combinatorial strategy (see Figure: "ISS workflow"), each probed gene is associated with a unique sequence of colors in a series of labeling/imaging/stripping cycles. By detecting the color sequence of the amplicons across cycles, we can deduce the identity of the underlying mRNAs.

**Intuitively, for this process to proceed smoothly, the images need to be perfectly aligned, so each amplicon can be uniquely identified in each cycle. Although this is ultimately achieved with computational tools, this task is much easier if the images are already reasonably aligned when the analysis starts. Please take exceptional care in mounting your samples consistently across cycles in the microscope.**

#### Glossary, useful definitions, abbreviations and clarifications:

**PLP**: Padlock probes

**FOV, or Tile:** Field of view, a specific area of an image

**RCP**: rolling circle product, or amplicon.

**Spot:** Fluorescent detection of an amplicon.

**Blob:** see "Spot"

**Detection event**: see "Spot"

**Anchor**: this is a deprecated staining method by which all the amplicons were labeled in a single color in the first round of imaging. This allowed generating a "map" of all the amplicons in the tissue as the first step of the analysis workflow. Currently this has been substituted by the "Pseudoanchor" (see below).

**Pseudoanchor**: This is essentially a computationally inferred anchor. All the amplicons in all the channels are merged together in a single image, so to draw a map of all them.

**Base**:  for historical reasons, we often refer to imaging cycles 1, 2 etc… and their outcomes, as "Base 1, Base 2, etc…". This is reflected in some of the file renaming steps, and occasionally in this manual. We acknowledge this might be confusing. Unless otherwise stated, a "Base" is the result of an imaging cycle in a given spot. The analogy is drawn from sequencing, in which a "base" is called by the detection of a fluorescence-emitting event. Likewise, a "base" in ISS is the fluorescent detection of the identity of an amplicon.

**Mipped**: maximum intensity projected image, i.e. taking the z stack of a given FOV, channel and cycle and projecting that into one single plane.

**Zero indexed**: python is zero indexed which means that it starts counting at 0, rather than 1. So if you for instance want to access the first element of the following list, i.e. the `str`: 'ISS':

```
my_list  = ['ISS','is','cool']
```

You would simply access it:

```
my_list[0]
Out[2]: 'ISS'
```

**Preprocessing**: the images from the microscope are taken in individual fields-of-view (FOV), which in our lab's microscopes have a size of 2048 by 2048 pixels with around 10-30 z planes. The FOVs have a 10% overlap between them. This means that we need to process the images by:

1. Projecting the z planes, resulting in 2D (2048x2048) images for each channel.
2. Aligning the 2D (2048x2048) images across channels.
3. Stitching the 2D (2048x2048) images into one large image for each channel.
4. Retiling the stitched images for each channel to reduce the computational demand of the subsequent steps.

**Decoding:** decoding refers to the process of deciphering the identity of the spots. This is based on looking at each single detection event and assigning the most intense channel for each cycle and thereby creating readout.

**Postprocessing:** this includes all the analysis that is done on the decoded spots. Dots are assigned to individual cells, cells are clustered according to expression similarity, etc…

#### Imaging and microscope requirements

Whichever microscope you use, it is important to reiterate the importance of **taking exceptional care in mounting your samples consistently across cycles in the microscope. Central to this process is the drawing of a ROI (region of interest) around the tissue region you want to image, using the microscope software. The ROI is typically specified in the first cycle of imaging, then saved and re-imported for the next cycles, so to ensure high quality alignment between the different images.**

#### How ISS analysis works

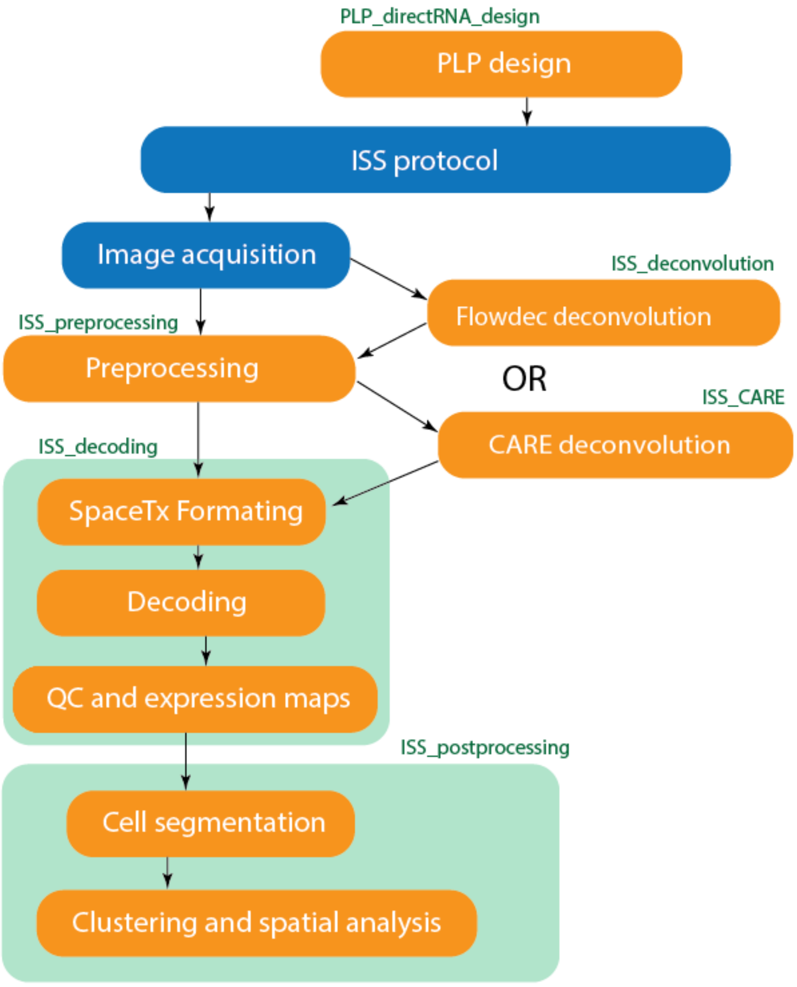

*Figure 2. Typical complete ISS image analysis workflow, showing optional modules that can be used to enhance image quality. Wet lab steps are represented in blue. Computational steps are represented in orange. Green represents the repositories from github.com/Moldia*

The scheme above illustrates the image analysis for ISS works. In the main processing pipeline, images are first preprocessed (organized, renamed, maximum projected, stitched, re-tiled, etc…). Once the preprocessing is complete, the images are transformed into SpaceTX format and decoded. The resulting datasets are stored as .csv, where each record corresponds to a dot, and columns are x,y coordinates, quality metrics, etc… Postprocessing of the decoded spots usually includes plotting the data, visualizing quality metrics, etc… as well as assigning spots to individual cells, and cluster the cells by expression similarities.

Two optional modules for image deconvolution are also provided. These modules feed into the main pipeline as depicted in the scheme above. Please refer to the "Advanced features" section in the manual.

### File naming and formats conventions

For image processing to proceed smoothly, it is highly recommended to follow the standard naming conventions used in our lab.

For microscopes brands that are Leica or Zeiss, softwares that export files in a different format, or users who prefer not to follow the easy path, the general rule is: find a way to export individual images (where individual image = a single tiff per tile, channel and z plane), together with a table including the xy positions of the tiles (this information is usually specified in the metadata associated with the images). The filename usually will include some flags to specify what is what, and you can try to adapt our preprocessing scripts to your needs.

Our file saving/handling rules are:
For Leica, we recommend the user to export, directly from the Leica software, the files as individual TIFFs. Each tiff file will represent a single plane of a single tile of a single channel, so thousands of individual TIFFs will be created in a typical experiment. It is important that the files are rigorously and consistently indexed, and this guaranteed by the following naming structure:
`TileScan 0--Stage01--C03.tif`
where `TileScan` represents the ROI, `Stage` represents the tile, `C` represents the channel, and the `--` sign acts as a separator (default on our Leica microscopes):

For Zeiss we give the users 2 options.
The first option is for the user to maximum-project the images manually inside the Zeiss software, and save the projections as tiffs, together with the xml metadata file.

The second option we recently developed, allows the user to input directly a czi file per cycle in the preprocessing step. The image data and metadata are automatically parsed from the CZI file and preprocessed automatically. Please refer to the "Preprocessing" section.

We are working on solutions for preprocessing Nikon and Olympus data and metadata. Please be patient while we do it. Otherwise you're very welcome to contribute and we can guide you through the steps to follow. Finally, if your microscope allows you to save the images in OMEtiff format, it might be possible to skip the initial step of preprocessing and proceed directly to stitching and registration, provided our file naming convention is respected.

#### Computer requirements

**The pipeline runs smoothly <u>on Linux computers.</u> Some people in our lab have managed to install and run it on Windows and MacOS, with a variable degree of frustration. We strongly recommend Linux.**

For the most basic usage of the pipeline (not doing any advanced image deconvolution or image restoration) any computer with at least 8 GB of RAM and i5 processor should be more than sufficient.

To run flowdec for image deconvolution (Czech et al. 2019) or CARE (Weigert et al. 2018) for image restoration, a CUDA-compatible GPU is needed. For flowdec deconvolution, In the lab we successfully used an NVIDIA Quadro RTX 4000, with 8 GB of RAM for small projects and an NVIDIA RTX A5000 with 24 GB of RAM for larger datasets. In the latter configuration, the speed is significantly higher. For CARE, the application of a model can be performed (at least in theory) on a CPU. This will be slower than using a GPU, but still reasonably fast. However, for training purposes we recommend our RTX4000 GPU configuration as the bare minimum, especially if working with large training sets.

ISS datasets can be very large, especially for large sections and/or high magnification acquisition. Furthermore, the data preprocessing steps essentially generate copies of the original files, renamed and reformatted, so the need for disk space quickly piles up. For easier handling of such datasets, we recommend saving the image data on SSD drives or fast on-line data storage systems. If possible, we suggest also to preprocess and decode the data using SSDs / online storage servers as output. This speeds up the data processing significantly.

### Basic introduction to installing python packages

If you are familiar with Python, Jupyter, etc… you can skip this section and go the the section covering the installation of the different analysis modules. If you are not, please read carefully. Before we proceed to explain how to install our ISS processing tools, we'll dig briefly into how python packages can be installed and how they are organized. There are three main ways to install a python package: using `conda`, using `pip,` and cloning repositories from github.

Using conda requires you to have **anaconda** installed, please see:
https://docs.anaconda.com/anaconda/install/

Using pip requires **pip** to be installed, please see:
https://pip.pypa.io/en/stable/installation/

Cloning of repositories requires having **git** installed, please see:
https://github.com/git-guides/install-git

To install the ISS processing packages, we will primarily make use of **git and the cloning of repositories from github**. The general syntax for using `conda`, `pip` and `git` is, respectively:

```
pip install package_name
conda install package_name
git clone https://github.com/package_name
```

### The jupyter project

We strongly believe in open and reproducible science, therefore, all of the github repositories are open for anyone to download and use. In addition, to make sure that the work that we do is reproducible, we make use of jupyter notebooks that can be shared among users. As you will see in the installation process, there is a file within the repositories with the extension `.ipynb`, these files are so called jupyter notebooks. These notebooks are essentially annotated code: blocks of codes that you can run interactively, with a (hopefully) clear explanation in plain english about what happens when you run each of them. Also, they contain a clear explanation of the inputs you have to provide and in which format.

#### jupyter lab vs jupyter notebook

jupyter lab and jupyter notebook are two ways to interact with notebooks. They are equally great and it comes down to personal preference. jupyter lab enables the user to have multiple notebooks in the same browser tab, whereas jupyter notebook opens one browser tab for each notebook that is open.

### Installation of the software toolboxes

The installation should be done using anaconda (Installing on Linux — Anaconda documentation) and separate conda environments (Managing environments — conda 4.13.0.post1+0adcd595 documentation), i.e. one conda environment for the preprocessing, one for decoding and one for the postprocessing, see down below how this is installed. We highly recommend creating and setting up the conda environment using the `.yml` file contained within the git repository, since this will ensure that all the package dependencies are installed correctly.

***Cloning the repository***

```
git clone https://github.com/Moldia/Lee_2023
cd Lee_2023
```

***Install the preprocessing module***

```
cd ISS_preprocessing
conda env create --name ISS_preprocessing --file preprocessing.yml python=3.7
conda activate ISS_preprocessing
python setup.py install
python -m ipykernel install --user --name=ISS_preprocessing
cd ..
```

***Install the decoding module***

```
cd ISS_decoding
conda env create --name ISS_decoding --file decoding.yml
conda activate ISS_decoding
python setup.py install
python -m ipykernel install --user --name=ISS_decoding
```

```
cd ..
```

***Install the postprocessing module***

```
cd ISS_postprocessing
conda env create --name ISS_postprocessing --file postprocessing.yml
conda activate ISS_postprocessing
pip install stardist
python setup.py install
python -m ipykernel install --user --name=ISS_postprocessing
cd ..
```

***Install the deconvolution module***

```
conda env create --file=ISS_deconvolution.yml --name=ISS_deconvolution python=3.6
conda activate ISS_deconvolution
python setup.py install
python -m ipykernel install --user --name=ISS_deconvolution
cd ..
```

***Install the CARE module***

```
cd ISS_CARE
conda env create --file=ISS_CARE.yml --name=ISS_CARE python=3.8.5
conda activate ISS_CARE
python setup.py install
python -m ipykernel install --user --name=ISS_CARE
```

Once these are installed, you can run `jupyter lab` and then toggle on the different environments.

### Run your first experiment:

#### Design probes for direct RNA detection.

##### Installation of the probe design software tools:

First of all you will have to install ClustalW2. Please follow the instructions at the following link: https://bioinformaticsreview.com/20210126/how-to-install-clustalw2-on-ubuntu/

From the cloned repository directory run:

```
cd PLP_directRNA_design
conda create --name probedesign python=3.9
conda activate probedesign
pip install cutadapt
python setup.py install
python -m ipykernel install --user --name=probedesign
```

if during the installation of cutadapt you get the error:

```
WARNING: The script cutadapt is installed in '/directory/blablabla/etc'
which is not on PATH.
```

Means you have to add the directory to PATH:
In linux this is done by typing:

```
PATH/directory/blablabla/etc:$PATH
```

This will need to be followed by the reloading of bash, done using the following line of code

```
source ~/.bashrc
```

To design probes for direct RNA detection, you need to input:

1. a reference transcriptome for the target organism, in FASTA format (.fasta/.fa/.fna). The transcriptome has to be downloaded locally. You can download most of the available transcriptome from NCBI or UCSC, or provide your own.
2. A list of genes for which you want to design probes, as in the example file. (https://github.com/Moldia/Lee_2023/tree/main/PLP_directRNA_design/examples/genelist_example.csv)
3. A gene-barcode association table, such as in: (https://github.com/Moldia/Lee_2023/tree/main/PLP_directRNA_design/examples/Lprobe_Ver2.csv)

The gene design pipeline should work for any species, provided you specify the correct gene names.

**However, one of the main functions of the probe design tool relies on a specific header format to extract the mRNA sequences based on the gene list provided. The function searches for the Gene names in the FASTA headers (line starting with >) using the format ().**

If your reference transcriptome doesn't specify the gene name in this format, the script will fail extracting the sequences, so in that case you will need to reformat the FASTA headers of your transcriptome. There are a number of tools and tricks to reformat fasta headers, but the instructions will really depend on how your initial headers look like. Consult a bioinformatician if you're completely lost.

The following link contains an example notebook with a detailed explanation of what's happening at each step, so we refer directly to the notebook as a "live manual" for probe design.

https://github.com/Moldia/Lee_2023/blob/main/PLP_directRNA_design/examples/PLP_design.ipynb

### ISS protocol and imaging

After ordering your padlock probes, it's time to run the ISS protocol in the lab. Despite this part will have a vast effect later on when processing and decoding the data, explaining this process is out of the scope of this manual. For further information about the protocol please refer to:

- **cDNA-based HybISS :** https://www.protocols.io/view/hybiss-hybridization-based-in-situ-sequencing-kqdg34357l25/v1
- **Direct RNA-based ISS:** https://www.protocols.io/view/home-made-direct-rna-detection-kqdg39w7zg25/v1

Please keep in mind that the probe design tool we refer to in this manual is only for direct RNA detection. You can find the old cDNA design scripts on our github.

### Preprocessing

#### Preprocess images from the microscope.

##### Objective of the preprocessing

The preprocessing module aims to convert files extracted from the microscope into a format suitable for decoding. Although the images coming out from a microscope typically cover a 3D space, our analysis works on 2D images. The first preprocessing step is therefore a maximum Z-projection of the raw images. The projected images ("tiles") are then stitched and aligned across imaging cycles. Finally, for computational reasons, the stitched aligned images will be resliced into smaller fragments. All this steps are contained in the repository found at www.github.com/Moldia/ISS_preprocessing

##### A note on intermediate files

Our preprocessing pipeline saves the output of all the intermediate preprocessing steps. This requires a lot of disk space, and feels often inconvenient. However, it can also help in troubleshooting when things go wrong, and to monitor rigorously each step, as you will learn in the next pages. In addition, this gives some flexibility, allowing the users to enter into the pipeline at any point.

In this step, the images are essentially read, organized, projected and rewritten in various formats, prior to the real image processing. **This is the entry point to the main analysis block: any user who wishes to analyze ISS images coming from microscopes that are not Leica or Zeiss, will likely have to intervene in these preprocessing routines.** Here's a summary of the main steps, so it's easier to understand what's going on under the hood and modify it if necessary.

In step 2 of the following routine, a `preprocessing` folder is created (in the folder the user specifies as output of the preprocessing functions). Several sub-folders will be created along the way, each one of them from one of the steps 2-5.

1. **Export images from the microscope software (this step happens in the microscope):**
    a. **Leica**: The data input will be the exported tiff files directly from the leica microscope. This will include the metadata as well but there is **no need to fiddle with the raw data** here. This is the default saving setup on the Nilsson's Leica microscopes, we recommend other users to strongly avoid .lif and .lof data formats from Leica and rather set their saving defaults to TIFF format.
    b. **Zeiss**: if the users desire to decode raw (non-deconvolved) images or apply CARE as a denoising method, they can maximum project and export the images on ZEN. However, this can be very tedious, so we now developed a function to automate this process, and also to assign our standard filenames to the output files. In this case, the user will just have to save the .czi files and apply a dedicated function in the next step. The function will extract and maximum-project the relevant files, and save them with the appropriate naming in the `/preprocessing/mipped/` folder, ordered by /`Base_n/` subfolders (see figures 3 and 4). The function will automatically extract the xy position of the tiles from the metadata contained in the .czi file and save this information in the `tile_coordinates.csv` file in each of the pertinent subfolders. Please check the paragraph related to the `process_czi` function
2. Once the images are **exported**, they will be **maximum projected** (unless they have been exported as already projected images, which is possible in ZEN), and saved in the `mipped` subfolder.
3. Following the projection, the images will be converted to OME tiffs files and saved into the `OME_tiffs` subfolder (see image below).
4. The **OME tiff** files will then be used for **alignment** and **stitching** using ASHLAR: Alignment by Simultaneous Harmonization of Layer/Adjacency Registration.
5. ASHLAR (Muhlich et al. 2022)will generate aligned and stitched images that are then **resliced** to reduce computational demands. The stiched images will be saved into the `stitched` subfolder, then resliced and saved one last time into the `ReslicedTiles` folder.

After the preprocessing step, your preprocessing folder should look as in the following image.

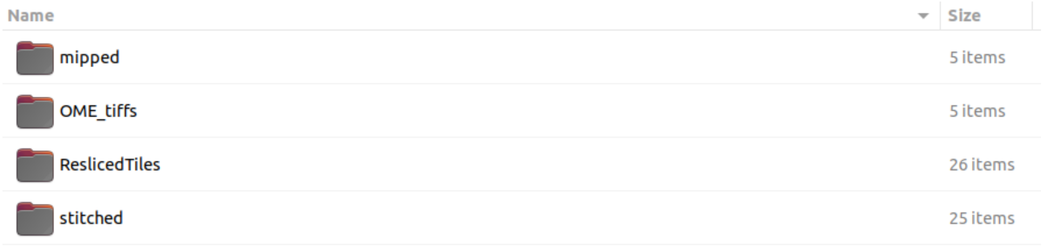


*Figure 3. Content of the `preprocessing` folder at the end of the preprocessing step*

We can peek inside each one of these folders, to have an idea of what to expect. The `mipped` subfolder should look like in the following figure (for an experiment of 5 ISS cycles). Each folder now contains the maximum projection of images for a given tile and channel, as well as the corresponding metadata. **Please note that in this step we use the nomenclature “Base” to indicate the imaging cycle, with an index starting in 1 for the first cycle.**

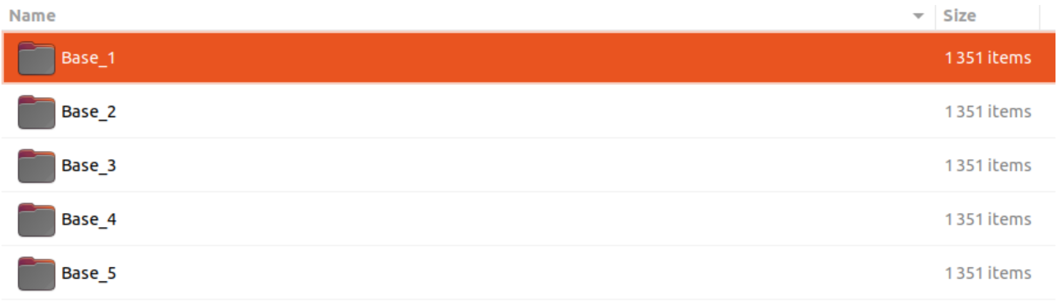


*Figure 4. Content of the `mipped` folder created during the preprocessing step*

Here's an example with some of the content of the `Base_1` folder.

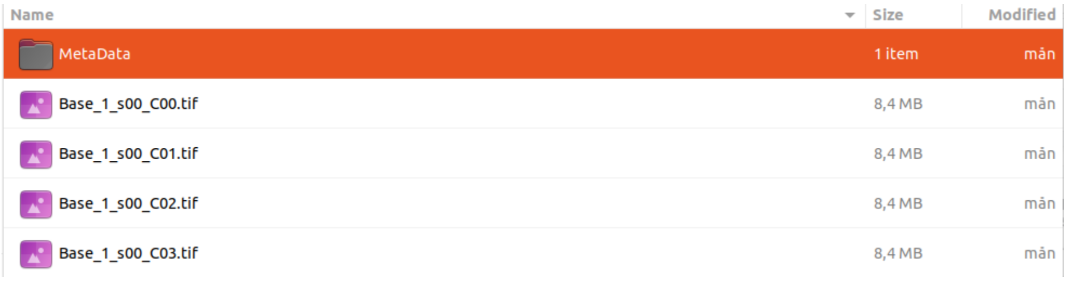


*Figure 5. Content of one of the `mipped` subfolders that are created during the preprocessing step*

The content of the `OME_tiffs` folder will look like this:

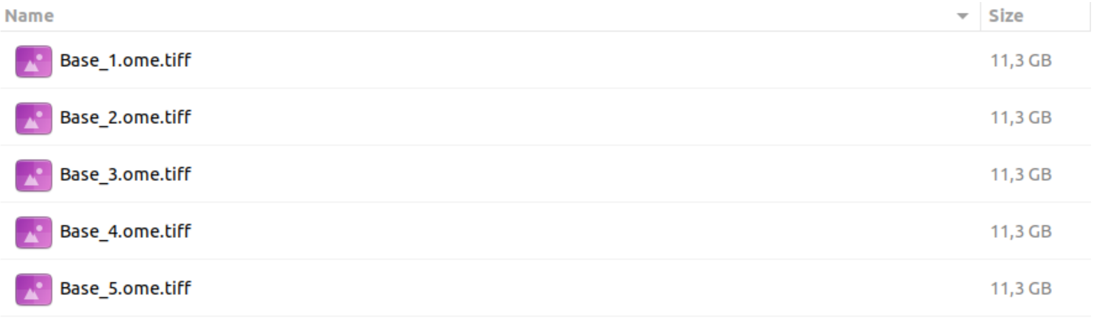


*Figure 6. Content of the `OME_tiffs` folder created during the preprocessing step*

**Please note that we keep the "Base" nomenclature in this step**. Now all the channels and tiles for a given imaging cycle are wrapped into a single file.

The next folder to be generated should be the `stitched` folder, and should look like this: each file is now a single stitched image, with a single cycle and channel. **Note that the**

**nomenclature to identify the imaging cycles has changed, and for this step we use "Round". Also note that Round starts with a value of 0.**

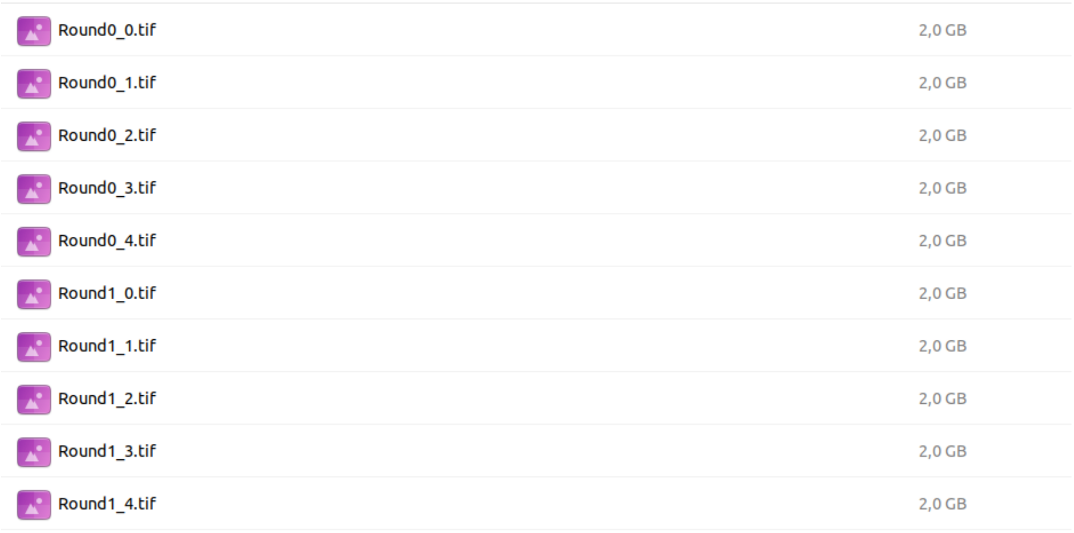


*Figure 7.Content of the `stitched` folder created during the preprocessing step*

The last folder to be created and populated is the `ReslicedTiles` subfolder, and should look like in the following picture. **Please note that the nomenclature to identify the imaging cycles has changed once again to "Base" starting with an index of 1.**

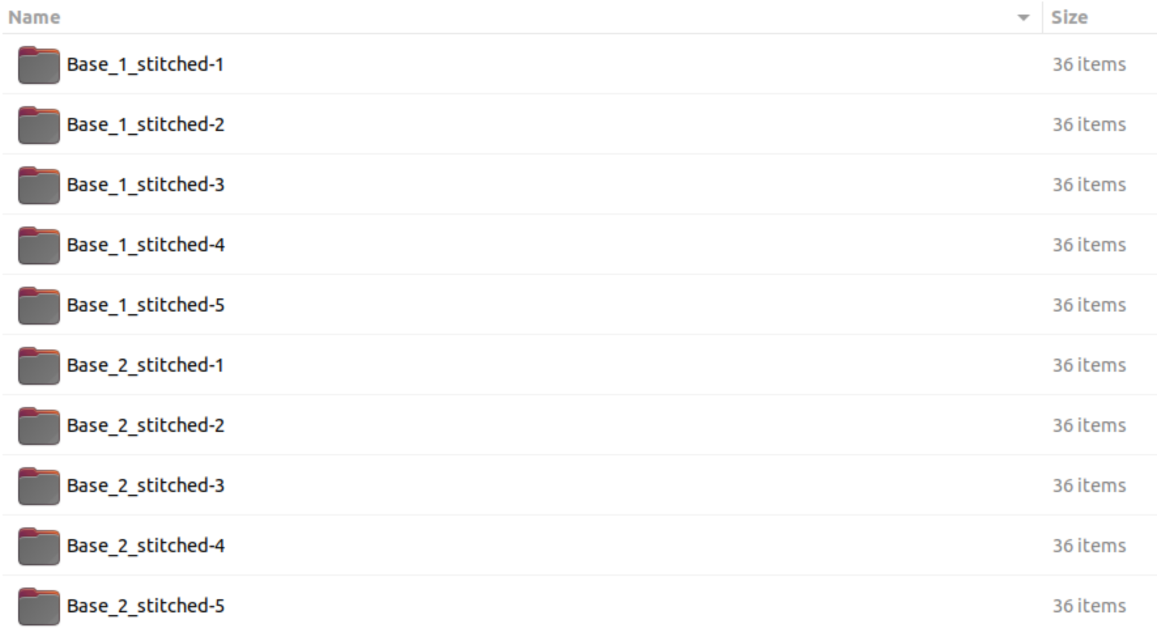


*Figure 8. Content of the `ReslicedTiles` folder created during the preprocessing step*

We acknowledge all of this might be confusing, and in a perfect case the user would never have to inspect what is going on under the hood. However, it is sometimes useful to know what to expect from the various preprocessing steps, so to figure out where the user got stuck in case of problems. These folders are created and populated in real time, so they are also a good way to monitor the progress of the analysis while it gets done.

#### Running preprocessing

When running the preprocessing, there are some things that need to be considered. First of all, what is your input data? Which microscope did you use? You can enter the preprocessing at any point, i.e. if you have aligned and stitched images, you can simply enter the workflow at the point of retiling and then proceed to the decoding. In addition, if your images are already projected into one plane, you can move directly onto generating the OME tiffs. Finally, if your microscope allows you to export projected OMEtiffs, you could perhaps start right away from registration and stitching.

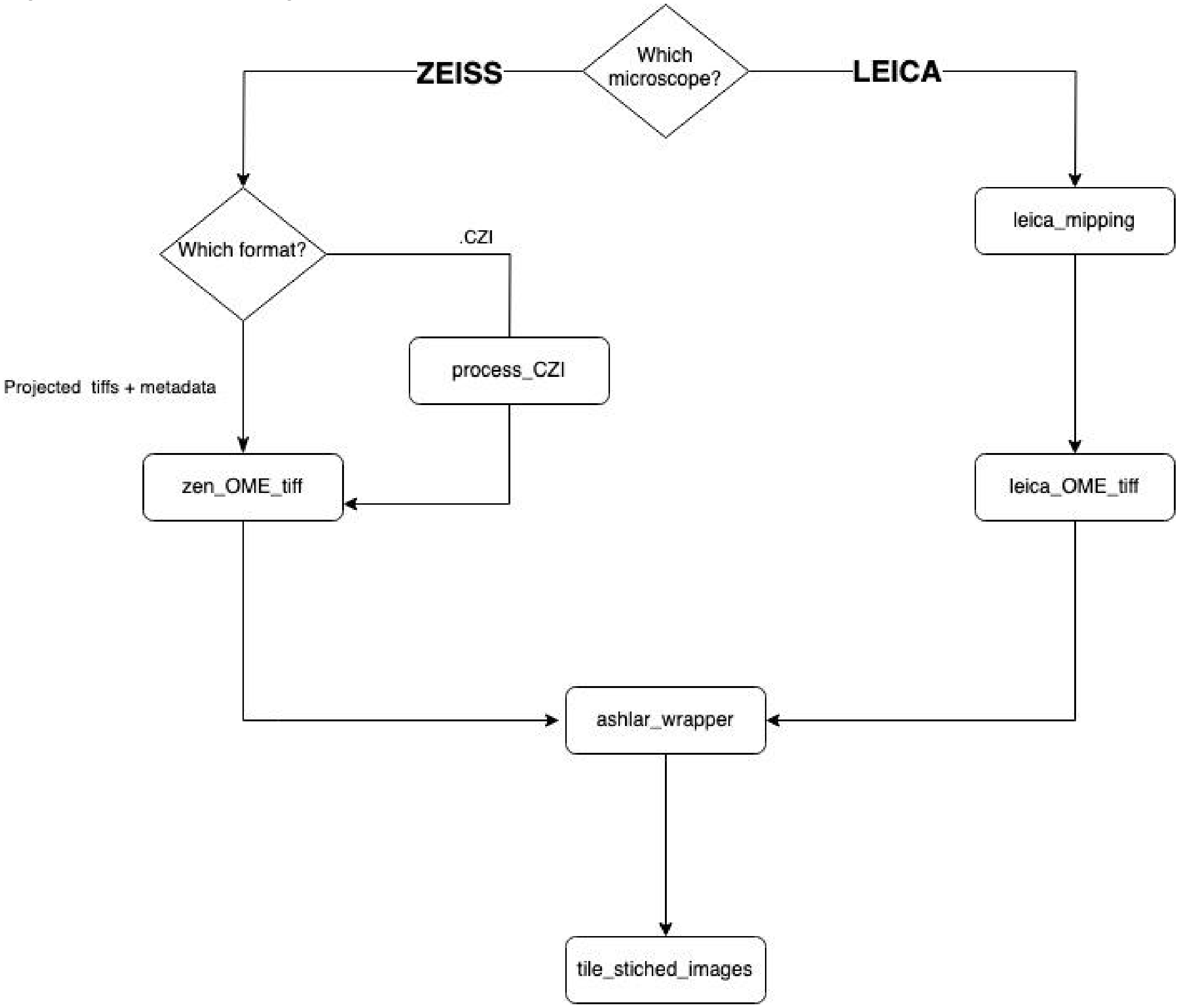


*Figure 9. Functions in the preprocessing module, together with a decision tree describing their use*

#### preprocessing_main_leica

This function (not depicted in the diagram above) wraps all the Leica related functions and runs them under the hood for simplified preprocessing, down to and including `tile_stitched_images`. This can be run using the function:

```
from ISS_processing.preprocessing import preprocessing_main_leica
preprocessing_main_leica(input_dirs = input_dirs,
                output_location = output_location,
                regions_to_process = 8,
                align_channel = 4,
                tile_dimension = 4000,
                mip = False)
```

##### Explanation of the variables

`input_dirs` = type: `list`. A list of `str` to where the raw data is. Example:

```
['/media/cml/Roadrunner/220609_fetal_spinalcord_cycle_1/2022_06_09_17_26_34--220609_fetal_spinalcord_cycle_1001/TileScan 1/',

'/media/cml/Roadrunner/220609_fetal_spinalcord_cycle_2/2022_06_09_13_04_07--220609_fetal_spinalcord_cycle_2001/TileScan 1/',

'/media/cml/Roadrunner/220608_fetal_spinalcord_cycle_3/2022_06_08_18_48_18--220608_fetal_spinalcord_cycle_3001/TileScan 1/',

'/media/cml/Roadrunner/220608_fetal_spinalcord_cycle_4/2022_06_08_14_14_38--220608_fetal_spinalcord_cycle_4001/TileScan 1/',

'/media/cml/Roadrunner/220608_fetal_spinalcord_cycle_5/2022_06_08_09_48_57--220608_fetal_spinalcord_cycle_5001/TileScan 1',

'/media/cml/Roadrunner/220607_fetal_spinalcord_cycle_6/2022_06_07_13_34_32--220607_fetal_spinalcord_cycle_6001/TileScan 1/']
```

`output_location` = type: `str`. this is the path where you want the output to be. Ideally, this should be associated with some type of unique sample identifier. Example:

```
output_location = '/media/cml/hfsc_processing_2/HFSC'
```

This will be the path where you want the preprocessing output to be saved. Ideally, this should be associated with some type of unique project identifier. The format of this variable is `str`. In case multiple regions are being processed, the funcion will behave differently depending whether a trailing slash `/` is included or not in `output_location`:

- if a trailing slash is added, then subfolders for each one of the scanned regions will be created as _R1, _R2, etc...
- if a trailing slash is omitted, then `output_location` will be updated at each region iteration, and each one of the scanned regions will end up in a different `output_location` named as `output_location_R1`,`output_location_R2`, etc..

`regions_to_process` = type: `int.` This refers to the number of regions that you have in one scan. In the example above, we had 8 regions. If you simply have one region, specify 1.

`align_channel` = type: `int.` This refers to the DAPI channel based on the order that your microscope acquires the images. In our set up, this is the fifth channel which in python would mean that we put 4 (since python is zero indexed: see glossary above for what it means).

`tile_dimensions` = type: `int.` This refers to the number of pixels that you want to tile your images into during the reslicing process. I.e. if you specify 4000, your resliced images will be of the shape 4000x4000 pixels.

`mip` = type: `bool.` This specifies whether or not you want to run the Maximum-projection step.

#### Running each of the steps separately

Instead of running the main function as outlined above. We can also simply run the functions by themselves. The input directories would be the same as above. The leica_mipping will handle the different regions automatically but the subsequent steps will have to be run for each region one by one.

```
from ISS_processing import preprocessing

leica_mipping(
    input_dirs=input_dirs,
    output_dir_prefix = output_location
)

path = '/media/cml/hfsc_processing_2/HFSC_1/'
# create leica OME_tiffs
leica_OME_tiff(
    directory_base = path+'/preprocessing/mipped/',
    output_directory = path+'/preprocessing/OME_tiffs/'
)

# align and stitch images
OME_tiffs = os.listdir(path+'/preprocessing/OME_tiffs/')
OME_tiffs = [path+'/preprocessing/OME_tiffs/' + sub for sub in OME_tiffs]

ashlar_wrapper(
    files = OME_tiffs,
    output = path+'/preprocessing/stitched/',
    align_channel=4
)

# retile stitched images
tile_stitched_images(
    image_path = path+'/preprocessing/stitched/',
```

```
    outpath = path+'/preprocessing/ReslicedTiles/',
    tile_dim=2000
)
```

For more information about how to use these functions, check the Jupyter notebook about the preprocessing of Leica files.

For Zeiss, we don't have a main wrapper function and the different functions need to be run one by one. The typical order by which you would need to run these functions is:

`process_czi, zen_OME_tiff, ashlar_wrapper, tile_stitched_images`

### `process_czi`

In a typical Zeiss experiment you will have 1 single CZI file per region per cycle. These files need to be processed individually, but of course feel free to wrap this function and the others in a loop for more automated processing. This function extracts the images, organises them, and create maximum projections that are exported with a naming convention fitting the downstream processing steps.

It also parses the metadata and converts them for downstream processing.

`process_czi` takes as inputs the following arguments:

`input_file`: the path to the CZI file that you want to preprocess, down to the czi file (included)

`outpath`: the folder where you want to save the maximum-projected images. Ideally this would be a `/mainoutputfolder/region/preprocessing/mipped/` folder structure, for consistency with our way of organising the data.

`cycle`: here you have to manually specify to which ISS cycle the images refer to. This is a `int` number, where 1 refers to cycle 1 and so on. If `cycle=0` the function will not work.

`tile_size_x` and `tile_size_y`: these refer to the size in pixel of your camera field of view. Most cameras are 2048x2048, so that's the default if you don't specify them, but adjust them if your camera has a different field of view size.

### zen_OME_tiff

This is the function that **takes the projected images across channels and wraps them into a single OMEtiff per imaging cycle**. This steps organises the files corresponding to each imaging cycles in a specific way within a single file and requires the parsing of a Metadata file to arrange correctly the images in xy space. As explained before, the input for this function is the specific sub-folder within `output_location` one wishes to process. **This function does not**

**allow to process multiple regions in one go, and needs to be run on individual regions manually**. From the previous function step, the mipped images will typically have this name format: `Base_1_c1m01_ORG.tif`

Where: `Base` refers to the cycle number, `c` refers to the channel number, `m` to the tile number. The `_` separator is used to parse the information about each image from the filename.

`Base_1_c1m01_ORG.tif` will be split into `Base_1` `c1` `m01`. Position 1 indicates the cycle, position 2 indicates the channel in this case.

The `zen_OME_tiff` accepts the following arguments:

`exported_directory`: this is the directory containing the maximum-projected images, however they were generated. They will serve as the input files

`output_directory`: this is the output directory that will contain the OMEtiff files created by the function

`channel_split`: this specifies where the channel number is indicated in the filename (default=2)

`cycle_split`: this specifies where the cycle number is indicated in the filename (default=1)

`num_channels`: this specifies how many channels (DAPI included) the images have (default=5)

Again, please refer to the notebook about preprocessing of CZI files for more details on the use of these functions.

### Decoding

#### Objective of the decoding

In this step, we make use of the **starfish** library to extract the information contained in our images. *starfish* is a Python library for processing images of image-based spatial transcriptomics. You can read more about it here: [https://spacetx-starfish.readthedocs.io/en/latest/](https://spacetx-starfish.readthedocs.io/en/latest/)

In the following steps, we first format our images to a starfish-compatible format (SpaceTx), and provide some information about our experiment design.
Once this is completed, we can proceed to the actual decoding of the data from the SpaceTx images.

#### Format images into SpaceTx format

The first thing that we need to do before we can start the actual decoding is to conform our images (the resliced tiles) to SpaceTx compatible data, also known as the SpaceTx format. To read more about the **SpaceTx format**, read the following: [GitHub - spacetx/sptx-format: Image data format specification for spaceTx](https://github.com/spacetx/sptx-format).

To format data, we use the ISS_decoding package and specifically the `SpaceTx_format` module, which can be found in: `ISS_decoding.SpaceTx_format.` This we can import in the following manner: `import ISS_decoding.SpaceTx_format as STX`

```
import ISS_decoding.SpaceTx_format as STX
STX.make_spacetx_format_zen(
   path = '/home/cml/Downloads/sample1/',
   codebook_csv = '/home/cml/Downloads/codebook_no_Plp1.csv',
   filenames=['Base_1_stitched', 'Base_2_stitched',
              'Base_3_stitched', 'Base_4_stitched',
              'Base_5_stitched','Base_6_stitched'],
   tile_dim=2000,
   pixelscale=0.1625,
   channels=["AF750", "Cy5", "Cy3", "AF488", "DAPI"],
   DO_decorators=["AF750", "Cy5", "Cy3", "AF488"],
   folder_spacetx='SpaceTX_format',
   nuclei_channel=5,
)
```

#### Explanation of the variables

`path` = type: `str.` this is the path to where your sample folder is located (given that you are following the nomenclature outlined above in ***File naming and formats conventions***)

`codebook_csv` = type: `str.` this is the path to your codebook, which is a comma separated values file with no header. The first column contains the gene name, while columns 2 to 6 (in case of a 6 cycle experiment) contain numbers representing the expected positive DO_decorator in each cycle for that gene.

***Example***:

```
codebook_hfsc_take2.csv
RNASE1,1,4,3,2,1,1
COL9A1,1,2,2,2,1,1
EPCAM,1,1,3,3,1,1
S100A9,2,3,4,1,1,1
CCBE1,2,4,1,3,1,1
RELN,2,1,1,1,1,1
NRXN1,2,2,3,3,1,1
CLDN5,4,4,2,2,1,1
GPC5,1,2,4,3,1,1
CD36,2,2,2,1,1,1
BMP4,2,4,3,1,1,1
BMP6,4,3,2,1,1,1
BMP7,3,3,1,1,1,1
CNPY1,3,1,3,1,1,1
DBX1,1,4,2,3,1,1
DLK1,4,1,2,3,1,1
JAG2,4,1,2,4,2,1
DLX2,2,2,2,2,3,3
HEY1,2,4,3,3,3,1
HGF,4,4,2,1,2,1
```

`filenames` = type: `list.` This list contains the names of the subfolders in the ReslicedTiles folder. Depending on how many cycles you have in your experiment, you will have to shorten it.

For example, if you only have 4 cycles, you should remove: `'Base_5_stitched','Base_6_stitched'`.

`tile_dim` = type: `int.` The dimensions of your ReslicedTiles. `Default = 2000`.

`pixelscale` = type: `int.` this is the size of the pixels, given in microns. This is needed if you want the coordinates of the spot to be output in microns, and will depend on your microscope settings. Set to 1 if you want the data to be on a coordinate scale. `Default = 0.1625`.

`channels` = type: `list.` The channel imaging order. `Default = ["AF750", "Cy5", "Cy3", "AF488", "DAPI"]`

`DO_decorators` = type: `list` this depends on how you associate the numbers in the codebook (ie 1,2,3,4) to a specific list of colors (DO_decorator) . In our lab, 1,2,3,4 correspond to `["AF750", "488", "Cy3", "Cy5"]. Default = ["AF750", "488", "cy3", "Cy5"]`. Users who will follow our barcode design and readout should not change this.

`folder_spacetx` = this is the name of the SpaceTx_folder. `Default = SpaceTX_format`.

`nuclei_channel` = type: `int.` This is the number of the channel that corresponds to your nuclei stained image. `Default = 5.`

As this step proceeds, a `SpaceTX_format` folder is created inside the output folder specified by the user. At the end of the process, its content should look like this picture. <u>Pay particular attention to the file `experiment.json`, which will be the input of the following step</u>.

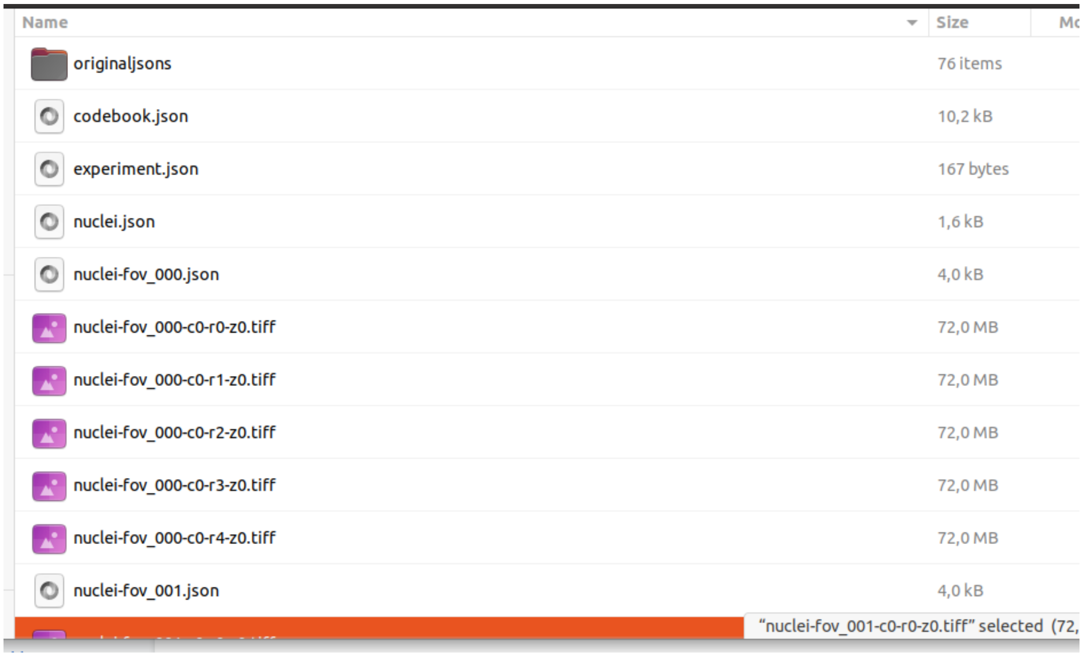


*Figure 10. Content of the* `SpaceTX_format` *folder created during the preprocessing step*

#### Extract and decode ISS data from the images.

Once the data has been converted into the SpaceTx format we can utilize the `process_experiment` function, which can be found in: **`ISS_decoding.decoding`**. This we can import in the following manner: **`import ISS_decoding.decoding as dec`**.

```
import ISS_decoding.decoding as DEC
DEC.process_experiment(
   exp_path = /home/cml/Downloads/sample1/SpaceTX_format/experiment.json',
   output = /home/cml/Downloads/sample1/starfish_output/',
   register = True,
   register_dapi = False,
   masking_radius = 15,
   threshold = 0.002,
   sigma_vals = [1, 10, 30], # min, max and number
   decode_mode = 'PRMC',
   normalization_method = 'MH'
)
```

#### What do we do when we run process_experiment?

The *process_experiment* function is run in order to decode the ISS experiment. The function will take every individual tile formatted in the SpaceTx format, detect every spot and assign to it the most probable code. Finally, the function will extract the XY position and identity of every spot that has been decoded in the tile. However, this process is actually a bit more complicated than this, and several steps happen within the process_experiment function. The main steps are as follows:

1. [OPTIONAL] Align/Register every tile based on either DAPI or a generated pseudo anchor.
2. Tophat transformation: Filters all images in all channels to enhance the signals and reduce the background, especially around the true signals, by applying a white tophat transformation
3. Normalization of the intensities: this is done to adjust the intensities between the channels and rounds. You can either chose MH or CPTZ.
4. Generation of a “pseudoanchor (see glossary)” based on the spots detected in all channels and all cycles. The spots will be detected on this pseudoanchor mask.
5. Detection of the spots using BlobDetector. We identify where our spots are using the pseudo anchor as a reference.
6. Decoding of the spots. Based on the xy(z) coordinates of the spots detected in the previous step, the intensity in all channels + all rounds is calculated. Based on this, the most probable barcode is calculated, associated with a gene, if possible, using the codebook and assigned a quality
7. Saving the results: the XY (Z) location of every decoded spot and its identity + quality is saved in a csv (for every tile).

All the steps are **repeated for every tile processed.**
Modifying some of the input parameters of the *process_experiment* function, it is possible to explore a range of settings to optimize the decoding for each experiment. These parameters are described in the following paragraph.

#### Parameters to modify when decoding (*process_experiment)*

There are many parameters that the user can change when decoding an ISS experiment. These parameters modify the behavior of some of the steps described above*:*

1. **Exp_path:** the user must specify the path where the experiment.json file is, within the SpaceTx_Format folder you have created when Formatting the images in SpaceTx Format.

2. **Output:** folder where the csv files containing the location of the decoded spots are saved. We recommend that this path points to a new folder, since we'll generate a csv for every tile processed.
3. **Register**: select *True* if you want to register/align your images and *False* if you don't want to align your images. This is recommended, as sometimes small adjustments in the registration of images are needed in this step.
4. **Register_dapi:** specify *True* if you want to align your images based on Dapi/ nuclear staining. If it's specified as *False,* the alignment will be done based on a pseudoanchor. This parameter only is considered if the parameter "Register is True".
5. **Masking radius:** the radius of your top hat filter. Depending on the size, it will "smoothen" your data to a different extent. The standard values go between 7 and 15. We'd recommend 7
6. **Threshold:** the most important parameter to tune. The threshold defines the minimum intensity that a spot should have in order to be detected. Decreasing the threshold will allow you to detect more spots, but if you decrease it too much you will start capturing background signals. It really depends on the experiment, but in general, using 0.002-0.005 should be fine. However, feel free to increase or decrease in an order of magnitude if you don't find your decoding accurate.
7. **Sigma vals:** correspond to min_sigma, max_sigma, num_sigma (explained in [Finding Spots with BlobDetector]). We don't use to modify them in the experiments, since they are already set to accurate values that allow us to capture signals with the size of our spots in 20X and 40X
8. **Decode_mode:** two options: 'PRMC': per round max channel OR 'MD', metric distance. After detecting the spots, the intensities of every channel and round of your experiment needs to be translated to a barcode, associated with a gene. For this, two strategies can be followed:
    a. PRMC (Per Round Max Channel): it calculates the most probable base on each cycle/round based on the channel with higher intensity. By combining the most probable bases on each cycle, it composes a code (i.e. 1232) and tries to look if this code is present in the codebook, assigning it a gene if present. If it's not present, it assigns it to non-assigned reads (NaN). The **advantage** of this approach is that it's an unbiased decoding strategy. The disadvantage is that, in bad quality experiments, we might call a lot of NaN. This is our preferred method for decoding
    b. MD (Metric Distance): metric distance tries to assign every spot detected to a code among the possible ones (among the ones specified in the codebook). This means that every spot will have a gene assigned to it. This is done by comparing the intensities in all channels present in the decoded spots with the expected signal generated by each of the genes in the codebook. As a result, we will also get a metric (distance) indicating how close each spot is to the nearest code present in the codebook. The lower the distance is, the better. The advantage of this method is that we can recover more spots than PRCM. The disadvantage is that we might overcall some of the fake signals and assign them to genes. Use this method with caution.

Whichever method, the output csv files consist essentially of the location (XY position) and identity of every spot decoded. However, keep in mind that not all the spots decoded might be accurate or good to use right away. Some of them might have a bad quality and, therefore, the data needs to be filtered. Refer to "[Understanding the results and quality metrics]" to understand quality scores and filtering methods.

9. **Normalization method:** ‘MH’ or ‘CPTZ’. The normalization of the images is done to adjust the intensities of all channels and rounds to facilitate the decoding. It can be done in two different ways:
    a. **MH**. **(Match Histograms)**. Matches the intensity histograms of all channels and rounds. It’s the preferred method.
    b. **CPTZ (Clip percentile to zero):** assumes that the differences in intensities come from outliers (super bright spots) or differences in the background. As a consequence, it just clips the values above a certain percentage (considered outliers) for every channel and round. This sometimes creates problems for tiles in the edge of the tissue, where the percentile of positive pixels can change compared to a tile in the middle of the tissue. Therefore, we might set the upper intensity limit in different values, causing potential tiling effects.

#### Combining all decoded tiles into a single file with the decoded spots

As we mentioned previously, every single tile is decoded independently. For this reason, the last step when decoding is to concatenate all the csv’s from individual tiles . For this, we use the following function.

```
import ISS_decoding.decoding as DEC
spots, spots_filt = DEC.concatenate_starfish_output(
   path = '/path/to/starfish/output/',
   outpath = sample
)
```

The *concatenate_starfish_output* function takes the “path” where your individual csv’ are and the output path (“outpath”), where the output should be saved. It concatenates all the tiles and saves the output in df_concat, a data frame including the spots assigned to NaN as well as the ones assigned to a gene from the codebook; and spots_filt, a data frame including the spots assigned only to a gene from the codebook.

#### Understand the results and quality metrics.

After combining the csv files for every individual tile, we should get as an output a big table (saved in csv) containing the information of our decoded spots. The main columns are 1. The location of the decoded spot (columns “xc” and “yc”) and 2. the identity of every spot (column: “target”). However, this dataset still contains a lot of spots that could potentially have bad quality. Therefore, we need to explore the quality of the signals.

##### How is the quality of the signals calculated?

###### What exactly is a "base" in ISS?

The following paragraphs use the concept of ISS "base". The concept is borrowed from SoLID or Illumina sequencing, in which a "base" is assigned (or "called") by the detection of a fluorescence-emitting event. Likewise, **we can define a "base" in ISS like "the true signal coming from an amplicon in a given cycle".** As in a sequencing, the base will have an associated quality score, depending on the signal to noise ratio (SNR).

To calculate the quality of a decoded spot, we need first to compute the quality of the so-called "base" for that spot in each cycle. This is essentially a SNR metrics: we measure the intensities in all channels, we assign the strongest channel as the "signal" and the remaining channels to "noise".

The quality of each base (Qbase) will be then described by the formula (signal/(noise+signal)), or in other words:

$$Qbase = \frac{Intensity\ Base\ max}{sum\ (intensity\ all\ bases)}$$

For each spot, we can then calculate two useful measures:
1) the quality_mean: the average of the Qbase across cycles. and
2) the quality_minimum: the lowest quality among all the cycles.
Other metrics that might give you information about the calling are:

- **Distance:** this is the distance (like hamming distance) from your called code to the closest code in the codebook. This is not a very interpretable measure, as it will depend on the size of your codebook. As a rule of thumb, the lower, the better. Read more in: https://spacetx-starfish.readthedocs.io/en/latest/gallery/how_to/metric_distance.html
- **Intensity:** mean intensity of your detected blob
- **Radius:** size (radius) of every detected blob

#### Plotting your quality metrics

There are several functions that allow you to explore how good your data is. In this section we describe the main functionalities that can be used to explore the quality of our dataset:

```
import ISS_decoding.qc_metrics as QC
```

**QC.plot_scores:** allows us to plot in the form of a histogram any quality parameter that we might have in our dataset. If decoding using "PRMC", where NaNs are detected, we can also plot the number of NaNs (non assigned) vs TRUE (assigned) reads against one of our quality scores. The function looks like this:

```
QC.plot_scores(reads,on='quality_mean',hue='assigned',log_scale=False)
```

This function needs as mandatory input the pandas dataframe (your csv containing the decoding output) [in this case, "*reads"]*. Then, several OPTIONAL parameters can be modified:

1. on: Quality metric (column name) that you want to plot as a histogram. We would typically like to test metrics like "quality_mean", "quality_minimum", "radius" or "intensity".
2. hue: **categorical** variable (column in the csv) that you want to use to split your columns in. The typical one would be "assigned", which, if you are decoding using PRMC, would allow you to visualize the percentage of assigned VS non assigned reads in different qualities. **NOTE THAT:** "assigned" is not a column in your "reads" object, and it will be created within the function based on whether the identity of every spot matches your decoding table or not.
3. Palette: specify the color palette of the plot based on the ones present in matplotlib
4. Log_scale: plot in log_scale if it's set to "True"

**How does the output look like?** We get two plots as an output: a histogram (left) and a density plot (right). They represent the same data, but in the density plot we have normalized the number of spots detected in every interval, so that the user can clearly observe the differences between your groups or, in this case, assigned vs non-assigned.

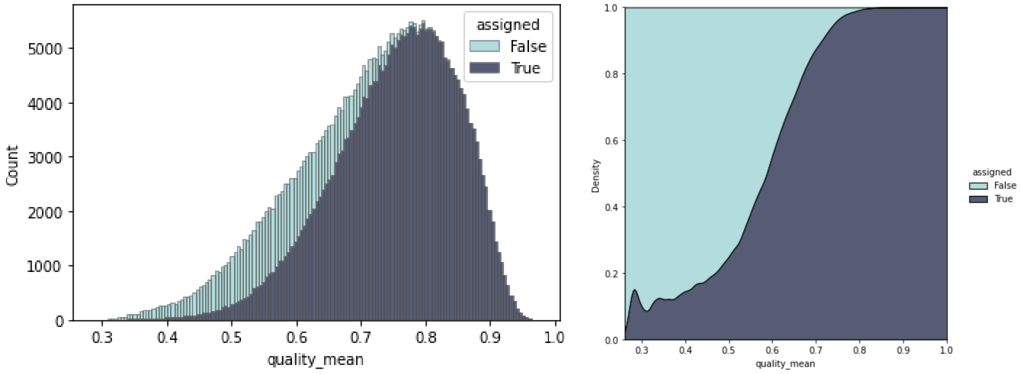


*Figure 11. Example output of the plot_scores function*

**Why should I use this for?** It can be useful to set a quality threshold. In the density plot above, the big shift between assigned VS unassigned occurs at quality_mean ~0.6 and, therefore, we would select this as our minimum_mean_quality.

**QC.compare_scores:** this function is used to plot more than one quality score at the same time. It represents the same as *plot_scores*, but for 2 quality metrics. It can facilitate the interpretation of how some metrics change. An example of use is:

```
QC.compare_scores(reads,score1='quality_mean',score2='quality_minimum',hue=
'assigned',kind='hist',color='#3266a8')
```

It needs as a mandatory input the pandas dataframe (your decoded .csv, in this example *reads)*. Also, the user needs to specify two scores (“score1” and “score2”), which are the columns containing the quality metrics that the user wants to plot.  Then, several OPTIONAL parameters can be modified:

1. Hue: **categorical** variable (column in the csv) that you want to use to split your columns in. The typical one would be “assigned”, which will split assigned vs non-assigned reads, as described above
2. Color: specify the color of the plot. It can be a color name or a HEX code
3. Kind: several types of plot can be done using this function. This include: “scatter” | “kde” | “hist” | “hex” | “reg” | “resid”. This options can be explored in: [https://seaborn.pydata.org/generated/seaborn.jointplot.html](https://seaborn.pydata.org/generated/seaborn.jointplot.html)

**How does the output look like?** The output is a 2D plot (depending on the parameter “kind specified”) that contains the relation of the two quality metric parameters evaluated, with the 1D-histograms of each individual axis on the margins of the plot.

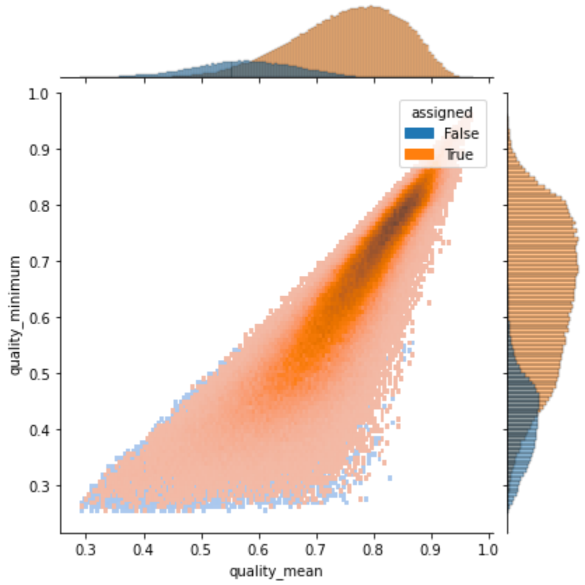


*Figure 12. Example output of the compare_scores function, where 2 different metrics are displayed on the xy axes.*

**Why should I use this for?** Comparing how one quality variable changes depending on another. This can clarify where to put your quality thresholds.

**QC.quality_per_cycle:** represents as a violin plot of the Qbase values obtained for the decoded spots on each cycle. It looks like:

```
QC.quality_per_cycle(reads,cycles=6)
```

*Table 3.*

It contains two mandatory parameters: the pandas dataframe with your results (a.k.a “reads”) and the number of cycles in your experiment (“cycles”). The output looks like:

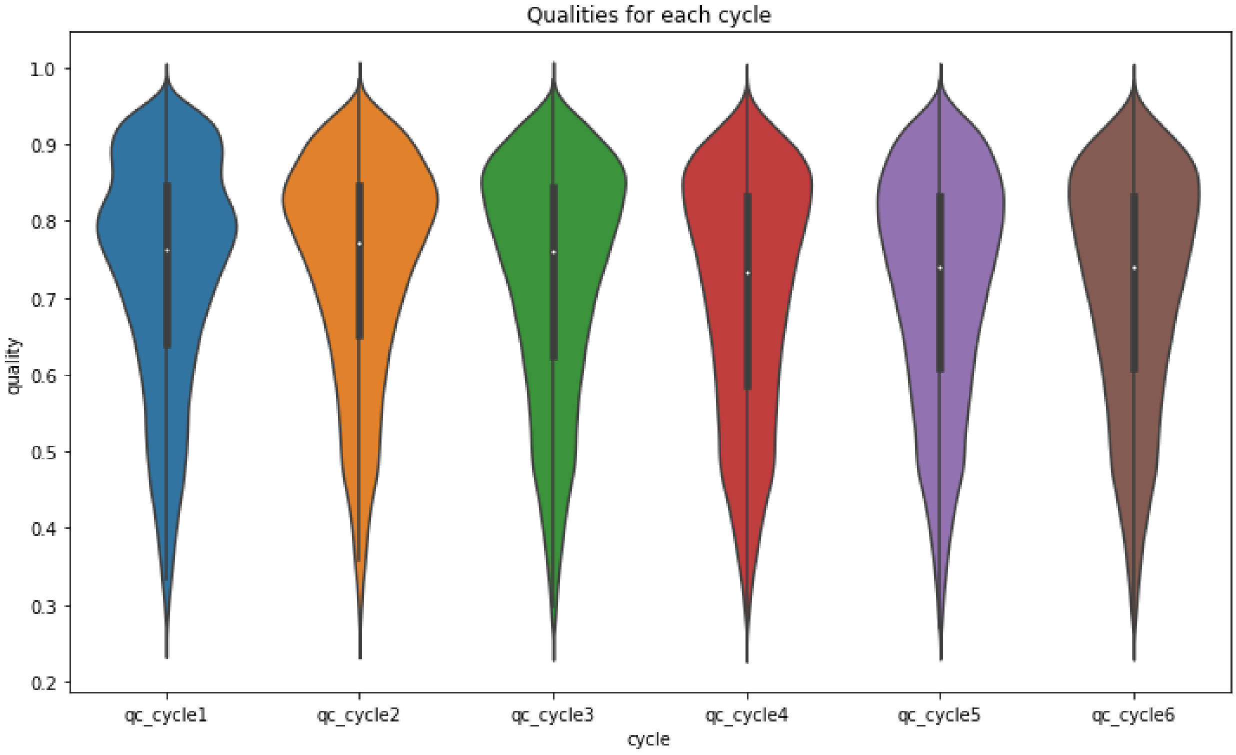


*Figure 13. Display of the quality of all the spots grouped by cycles.*

**Why should I use this for?** Checking if the quality of your spots is consistent throughout the cycles can help you spot cycles with lower quality. Low quality cycles typically bias or impair the decoding. A cycle with a low quality can be caused by different reasons including (1.) Bad alignment of the images for that specific cycle, (2). Bad detection of the signals (3). Out of focus images. There might be other causes, but they are usuallt unlikely. Overall, it points out the necessity of closely inspecting the raw images, and perhaps reprocessing or reimaging one of the cycles.

**QC.quality_per_gene:** this function is designed to evaluate the quality of all the reads assigned to each gene. Since every gene has an associated barcode with a specific sequence of colors across cycles, we might have a big quality bias between different genes just because of the barcode composition. The function looks like:

```
QC.quality_per_gene(reads,on='quality_minimum',gene_name='target')
```

It needs 2 mandatory parameters: your dataframe/csv with the decoded spots (reads) and “on”, where you need to specify the quality metric/ column that you want to evaluate. You can also add a gene_name column in case the identity of every spot is saved in a special column, but the default is “target” which is where we have the identity of our spots.  The output looks like this:

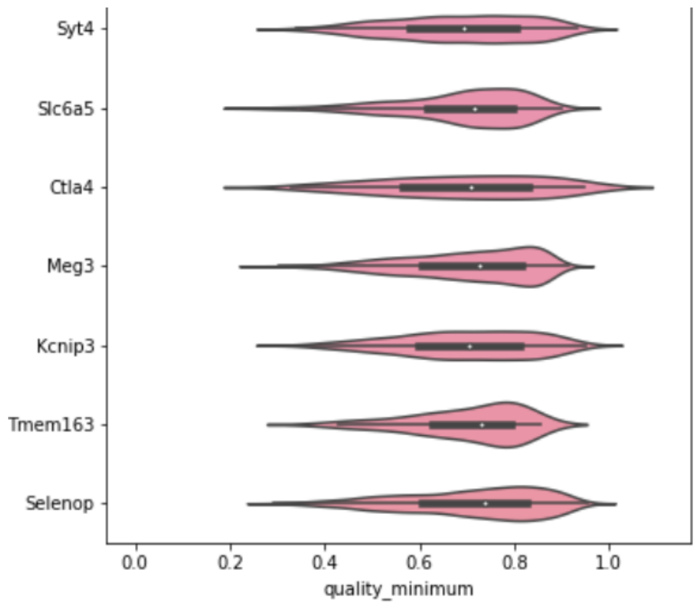


*Figure 14. Display of a quality measure for all the spots assigned to each gene*

You can see how for different genes the violin plot looks slightly different. This is important because, if you set the quality thresholds too high, you might lose the signals of specific genes and this can bias the results and interpretation.
**Output:** there is an optional output, which is a data frame with the mean quality of every gene.

**QC.plot_frequencies:** it can be used to plot the frequencies of every gene detected as an histogram. The code is:

```
QC.plot_frequencies(reads,on='target')
```

It needs as an **input** two different parameters: (1) your csv/dataframe (a.k.a reads in here) with the information of your decoding spots and (2) “on”: the column that contains the identity of every spot. As an optional output it can return the counts for every gene as a pandas data frame.

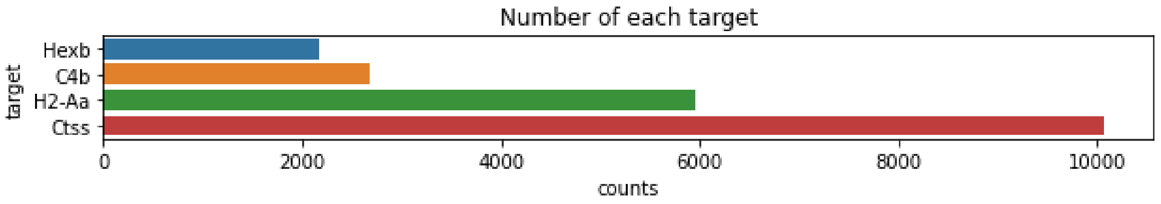


#### Filtering the results

Based on all the quality plots you generated in the last section, at this point you should have an idea about how to filter your data and whether it has a good quality. To filter it, we will use the function *filter_reads*.
Note about quality parameters: if you look back at how the Qbase measure is calculated, you will see that its minimum possible value is 0.25 for a 4 color decoding, or 0.2 for a 5 color decoding..

```
good_reads=QC.filter_reads(reads,min_quality_mean=0.6)
```

Using this function, we can input, as usual, our "*reads*" dataframe, containing the information about the decoded spots in a csv. Then, several optional parameters can be selected depending on how you want to filter the data:

- Min_quality_mean: minimum quality_mean that the reads should have
- Min_quality_minimum: minimum quality_minimum that the reads should have
- Max_distance: maximum distance that your decoded spots should have (only applies if metric distance decoding is used)
- Max_radius: maximum radius allowed. Everything above will be filtered out
- Min_radius: minimum radius allowed. Everything below will be filtered out
- Min_intensity: minimum intensity that the decoded spots should have
- Max_intensity: maximum intensity that the decoded spots should have

Usually a conservative method for filtering is to set min_quality_minimum=0.5 and discard all the reads that do not meet this criterion.

The **output** of this function (here named as good_reads, but you can give it whatever name you want), is a data frame containing the filtered dataframe. This can be saved as a csv and used as an input for visualization or downstream analysis.

#### Plot and visualize the decoded data.

Whether before or after filtering, the decoded data can be plotted using the function:

```
QC.plot_expression(reads_filt,key='target',colorcode=['red'],xcolumn='xc',ycolumn='yc',genes='individual',size=2,background='black',title_color='white',figuresize=(10,10),save=None,format='pdf')
```

The mandatory arguments of this functions are 1) a dataframe containing your reads, 2) key= the header of the column in the dataframe containing the gene names, 3) xcolumn and ycolumn, the column in the dataframe containing the dots' coordinates. Other arguments are:
genes = 'individual', plots one gene at the time, 'all' plots all the genes in a single plot
size= sets the dot size
background= sets the background color
title_color= sets the color of the title for each plot.
colorcode= sets the color of the expression dots

Another good option to explore ISS data in a more interactive way, make figures, etc… is TissUUmaps, from Carolina Whalby's team: https://tissuumaps.github.io/
We refer to the TissUUmaps manual for more information.

### Postprocessing

#### Objective of the postprocessing

The postprocessing toolbox drives the user through a set of steps for higher-level analysis. While some ISS users might simply be interested in analyzing the expression patterns of large number of genes, the full power of ISS and other multiplexed in-situ methods is best deployed when we can use the gene expression information as the basis for advanced quantitative analysis.
The postprocessing steps might involve assigning expression spots to individual cells, clustering the cells by similarity, integrating the ISS datasets with prior scRNAseq datasets, etc…

#### Cell segmentation

**Cell segmentation is the task of splitting an image into smaller blocks, where each block corresponds to the space occupied by a cell or a nucleus**, such as in example below.

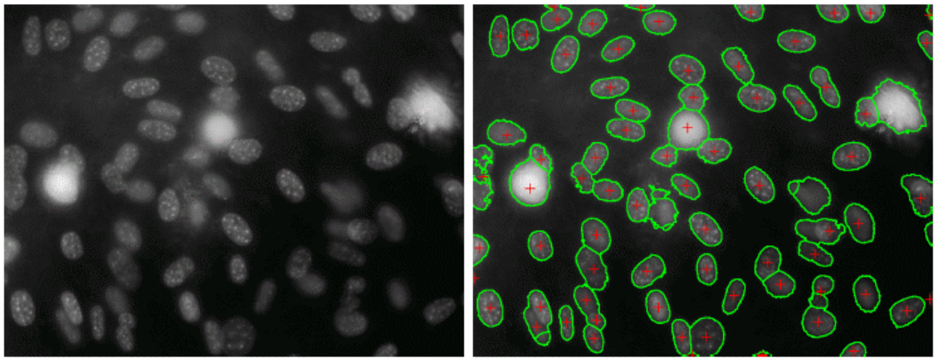

*Figure 15. Example of nuclei segmentation from Cellpose manual.*

By extension, **cell segmentation allows us to assign each ISS spot to an individual cell**. This will allow us to create, for each individual cell, a quantitative description of its mRNA expression profile. This is a necessary step to answer questions such as "how many types of cells do I see in a tissue?", "how many cells of each type compose the tissue?", "how do different cell types compare to each other?", etc…

##### Nuclei based cell segmentation

In the lab, we use primarily two strategies for nuclei based segmentation. Both are based on machine learning models.

###### cellpose ([GitHub - MouseLand/cellpose: a generalist algorithm for cellular segmentation with human-in-the-loop capabilities](GitHub - MouseLand/cellpose: a generalist algorithm for cellular segmentation with human-in-the-loop capabilities))

Cellpose (Stringer et al. 2021)has a user interface that can be used to train machine learning models, and more importantly, you can use it to evaluate already existing models on your dapi stained images.

```python
import ISS_postprocessing
ISS_postprocessing.segmentation.segment_tile(
    sample_folder,
    segment = True,
    dapi_channel = 5,
    diam = 40,
    expand_tile = False,
    expanded_distance = 20,
    big_section = False,
    output_file_name='cellpose_segmentation.npz',
)
```

##### Explanation of the variables

`sample` = type: `str`. This is the path to your sample folder.

`segment` = type: `bool`. If you want to segment or not. You should put False if you want to restitch already segmented tiles. `Default = True`.

`dapi_channel` = type: `int`. The imaging channel. `Default = 5`

`diam` = type: `int`. The diameter that cellpose will use to look for cells. `Default = 40`.

`expand_tile` = type: `bool`. Whether to expand the cells or not. `Default = True`.

`expanded_distance` = type: `int`. The number of pixels to expand the cells. `Default = 30`.

`big_seciton` = type: `bool`. If the section is very big you might need to split the output segmentation mask into two. If you select `True` here the output will be split into two. `Default = False`.

`output_file_name` = type: `str`. The name of the output file. `Default = cellpose_segmentation.npz`.

##### stardist ([StarDist - Object Detection with Star-convex Shapes]())

In general I find the pretrained models in stardist ((Schmidt et al., 2018) to be better than cellpose and it runs way faster. It is also a bit more straightforward to run since it allows you to input a full dapi image that can then be split into however many tiles you want.

```python
import ISS_postprocessing
ISS_postprocessing.segmentation.stardist_segmentation(
    image_path = '/path/image.tif',
    output_path = '/path/',
    model_name = '2D_versatile_fluo',
    n_tiles = (4,4),
    expand_cells = True,
```

```
    expanded_distance = 20,
)
```

This will output a coo matrix saved as a npz file. This is to ensure that the matrix does not take up too much space. Additionally, this is the format that is needed to run `pciSeq`.

##### Explanation of the variables

`image_path` = type: `str`. The path to your stitched dapi image.

`output_path` = type: `str`. The path to where you want the segmentation file to be stored.

`model_name` = type: `str`. The name of the model that you want to use. `Default = '2D_versatile_fluo'`

`n_tiles` = type: `tuple`. The dimensions of that you want the segmentation to be split up into. `Default =(4,4).`

`expand_cells` = type: `bool`. Whether to expand the cells or not. `Default = True.`

`expanded_distance` = type: `int`. The number of pixels to expand the cells. `Default = 20.`

#### Pixel based classification

There are also great alternatives when the nuclei are hard to segment using either of the approaches above. One such option is to use the pixel classification function in ilastik. To learn more about ilastik and the pixel classification function, visit: [ilastik - Pixel Classification]. What this essentially enables you to do is to draw on top of your dapi stained images to determine whether or not a pixel belongs to a cell or not. This is fast and works great on samples where we can not use cellpose or stardist.

#### Spot based segmentation

There are also other spot based segmentation strategies that one can use to segment data, one of these that we find to be of particular good use is Baysor (GitHub - kharchenkolab/Baysor: Bayesian Segmentation of Spatial Transcriptomics Data). What is neat about baysor is that it doesn't rely on nuclei staining for segmentation of cells. Some tissues can be particularly hard to segment due to very dense nuclei or nuclei with ‘hazy’ outlines, developmental tissues in particular. In addition, what makes baysor such a useful tool is that it can leverage previously segmented cells from stardist or cellpose, and put different confidence levels on the previously segmented cells.

Follow the steps as outlined in the link above to install. This requires dense data to work well but under the right circumstances, can perform well for ISS data.

### What can we do downstream of the cell segmentation?

For an example notebook on the downstream analysis of ISS data, see https://github.com/Moldia/ISS_postprocessing/blob/main/ISS_postprocessing_tutorial_scanpy.ipynb. This notebook includes the segmentation of the cells using stardist, creating the anndata objects and the leiden clustering of the cells.

#### Probabilistic cell typing by in situ sequencing (pciSeq)

Probabilistic cell typing by in situ sequencing was published in 2020 (Qian et al. 2020)and it is a way to probabilistically assign genes to cells and cells to cell type based on a single cell RNA sequencing reference dataset. This can be found under `ISS_postprocessing.ISS_postprocessing.pciseq.` There is a great example of how to run this: [GitHub - acycliq/pciSeq: A probabilistic cell typing algorithm for spatial transcriptomics]() and we refer to the documentation outlined here.
You can find a guided tutorial on pciSeq in our github.

#### tangram

tangram (Biancalani et al. 2021) enables mapping single-cell gene expression data onto spatial gene expression data. We refer to the documentation on [GitHub - broadinstitute/Tangram: Spatial alignment of single cell transcriptomic data]().

#### Format ISS data as Annotated data objects

Given the vast amount of tools that revolve around the anndata format (annotated data: [AnnData]()), we might want to convert our ISS data into this format. This will allow us to process our data in [Scanpy](), as if they were single-cell data.

The first thing that we need to do is to assign the reads that we get from the `ISS_decoding` module to the segmented cells from the **`ISS_postprocessing`** module. The function to create scanpy compatible files is stored in the **`ISS_postprocessing`** module.

##### Create annotated data objects

```
adsp = ISS_postprocessing.annotated_objects.create_anndata_obj(
        spots_file = "decoded_aligned_MD.csv",
        segmentation_mask = "/path/to/coo.npz",
        output_file = "/path/to/annData.h5ad",
        filter_data=True,
        metric = 'distance',
        write_h5ad = True,
        value=  0.4,
        convert_coords = True,
        conversion_factor = 0.1625
)
```

**`ISS_postprocessing.annotated_objects.create_anndata_obj`**. Depending on the size of your tissue and the capacity of your machine, this can either be done on a tile-by-tile basis or the whole section at once. We prefer the latter since this removes the issue of spots and cells located on the edges of the tile, being assigned wrongly.

Explanation of the variables

`spots_file` = type: `str`. this is the path to your spots file.

`segmentation_mask` = type: `str`. The path to the coo file generated from the segmentation.

`output_file` = type: `str`. Path to the output file.

`filter_data` = type: `bool`. True or False, to filter the data or not. This option should be used only if inputting the raw reads (unfiltered) or additional filtering is needed.

`metric` = type: `str`. The metric we want to filter the data based on, if `filter_data=True`.

`value` = type: `int`.value used for filtering, if `filter_data=True..`

`write_h5ad` = type: `bool`. True or False, should the annData object be written to a h5ad file.

`convert_coords` = type: `bool`. True or False, should the coordinates be converted from microns to pixels.

`conversion_factor` = type: `int`. The conversion factor to go from microns to pixels.

### Cluster your cells

#### Purpose of clustering

The main purpose of clustering is to detect the main cell populations present in your dataset. With *de novo clustering* you will identify the main clusters present in your dataset using **only** the expression information about the genes included in your panel. This means the variability between cells needs to be captured in the probe panel you used.

#### When should you do it?

Clustering cells (*de novo*) can be done when you have enough reads/cells in your dataset. This means that if you have (on average) fewer than 5-10 reads/cell, you won't have enough information to cluster your data. Any number of genes will work, provided that they capture the diversity present in your cells. As a rule of thumb, more markers and more reads will produce more reliable clustering. If your purpose is to map your ISS data on scRNAseq clusters, alternatives such as pciSeq / Tangram might be better than using *de novo* clustering.

#### Procedure

This analysis is based on the popular Python packages Scanpy and Squidpy. You can read further (documentation in https://scanpy.readthedocs.io/en/stable/index.html and https://squidpy.readthedocs.io/en/stable/ respectively).

##### Formatting the data

As a first step we import the data containing the information of every cell in a cellXgene table, stored as AnnData object (documentation in https://anndata.readthedocs.io/en/latest/). For this, we need to first read our data, using pandas like follows:

```
dataset=pd.read_csv(<PATH TO YOUR CSV>)
```

Typically, your dataset will be a table in which every row is a cell, and the columns describe the expression of every gene. Some columns are reserved for the metadata (everything that is not strictly gene expression). This metadata includes things such as: the size of the cell, the XY positions of the cell or other information.
We need to split the dataset into (1) expression data, so every column contains the expression of a gene and (2) metadata. For this, we need to explore where each of our columns belong to. In this case, we create two needed dataframes, one for expression and the other for metadata. In this example, columns 0 to 80 correspond to metadata and 80 to 220 correspond to expression:

```
metadata=dataset.iloc[:,0:80]
expdata=dataset.iloc[:,range(80,220)]
```

Then, we'll need to create the main object that we will further use, and that we do it by using the following command:

```
adata = sc.AnnData(expdata)
adata.obs=metadata
```

The object containing our data is called *anndata* (type of dataset). In the scripts, we usually call it *adata,* but one can give it any name.This type of objects have always three main "subobjects", .X, .var and .obs As a summary:

- The expression of your cells is stored in **adata.X**. This is a matrix with the dimensions being number of cells by the number of genes
- The metadata of every cell  is stored in **adata.obs**. In here we have the additional information we have about every cell. In spatial methods, here we can have the position of our cells (XY position), the number of reads in every cell, as well as the cell type label, etc… It's a matrix/data frame with dimensions being the number of cells by the number of metadata terms.
- The metadata associated with every gene  is stored in **adata.var**. This contains the name of every gene, but then we can add other information about each gene, such as if it's a highly expressed gene, or their GeneID. Its dimensions are Number of genes by number of metadata.

You can see in this plot the structure of an AnnData object:

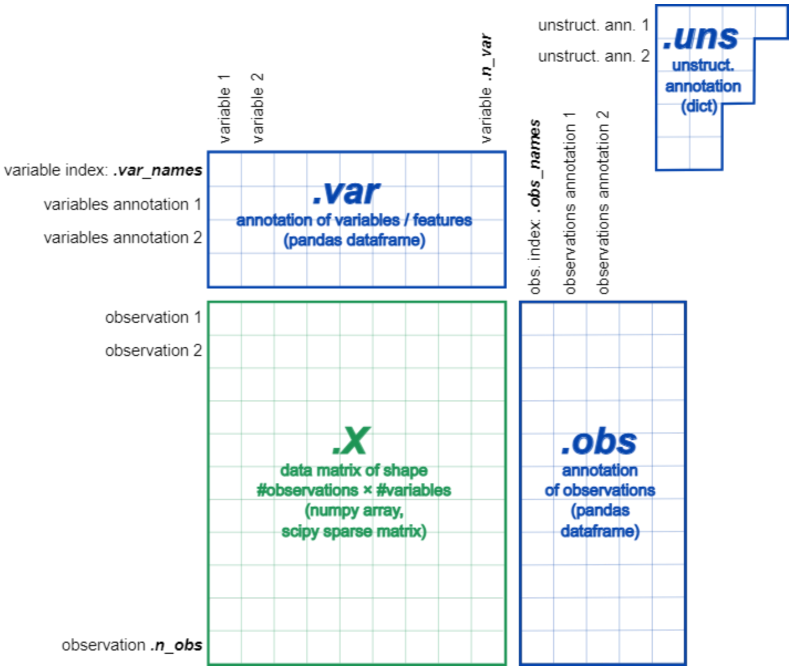


As you can see, your central matrix is your expression matrix, which is a cell by gene matrix (stored in adata.X).Then both .var and .obs share one of their dimensions with .X, matching with either the number of genes or number of cells respectively. An additional place where to store information about this object is adata.uns, which corresponds to unstructured data. This refers to information that's not related with the number of genes or the number of cells, for example, the UMAP coordinates or the differentially expressed genes in every cluster.

In practice, if you just type the name of your object of your script this is what you get:

```
adata
```

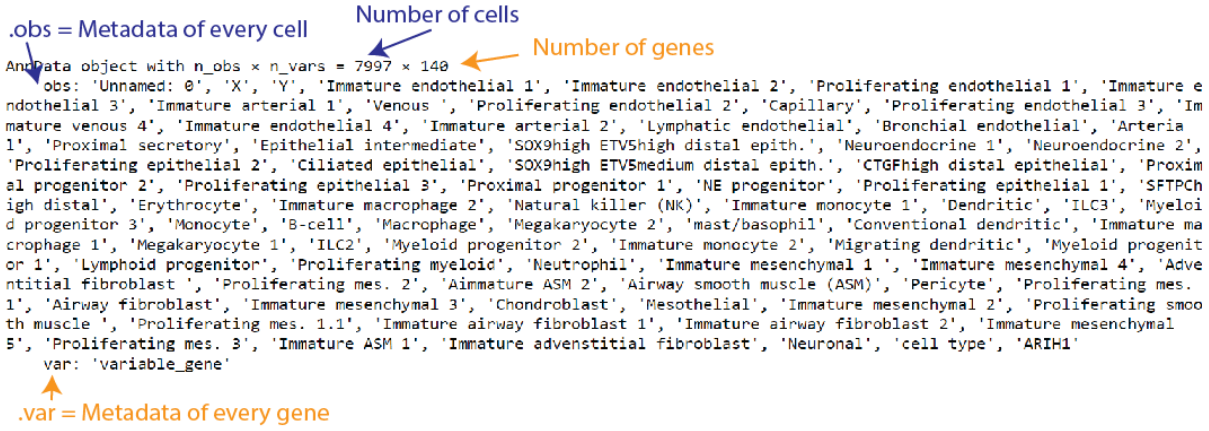


For exploration in order to access each of these terms, we can just write the name of our anndata object (in this case adata) and any of these terms.

#### adata.obs

Adata.obs returns a dataframe with the cell-related metadata:

| | Unnamed: 0 | X | Y | Immature endothelial 1 | Immature endothelial 2 | Proliferating endothelial 1 | Immature endothelial 3 | Immature arterial 1 | Venous | Proliferating endothelial 2 | ... | Proliferating mes. 1.1 | Immature airway fibroblast 1 |
|---|---|---|---|---|---|---|---|---|---|---|---|---|---|
| **0** | 0 | 988.532325 | 3787.312606 | 4.229731e-13 | 1.217327e-10 | 2.910561e-11 | 1.690861e-14 | 5.669160e-12 | 4.048466e-08 | 4.986064e-07 | ... | 2.483804e-11 | 4.086888e-11 |
| **1** | 1 | 1069.522775 | 3984.402494 | 7.688518e-06 | 1.076019e-05 | 1.024221e-05 | 5.217759e-06 | 3.937803e-06 | 4.062558e-05 | 7.545922e-06 | ... | 5.829106e-05 | 6.948951e-04 |
| **2** | 2 | 1014.118412 | 4170.700890 | 4.681991e-02 | 4.147485e-02 | 4.706028e-02 | 4.887775e-02 | 2.672377e-02 | 5.769456e-02 | 2.077321e-02 | ... | 8.316057e-04 | 7.012579e-04 |
| **3** | 3 | 1085.879776 | 4122.941906 | 1.435361e-02 | 8.028734e-02 | 4.917569e-02 | 1.310387e-02 | 2.058924e-01 | 4.113200e-02 | 2.010187e-01 | ... | 7.056170e-09 | 6.422320e-10 |
| **4** | 4 | 1093.771630 | 3619.763833 | 2.516980e-28 | 2.425292e-25 | 1.312165e-27 | 6.754184e-29 | 5.022944e-28 | 4.158016e-18 | 4.536668e-24 | ... | 1.028110e-12 | 1.846184e-13 |
| **...** | ... | ... | ... | ... | ... | ... | ... | ... | ... | ... | ... | ... | ... |
| **7992** | 7992 | 12373.000000 | 5095.000000 | 2.003389e-04 | 2.439488e-04 | 1.397369e-04 | 2.648583e-04 | 2.425629e-04 | 1.288632e-04 | 3.400722e-04 | ... | 4.484618e-02 | 5.860223e-02 |
| **7993** | 7993 | 12385.089530 | 6011.623672 | 2.179862e-05 | 3.217089e-05 | 1.111304e-05 | 2.391398e-05 | 1.889256e-05 | 1.990930e-04 | 2.772565e-05 | ... | 5.577385e-03 | 7.508577e-03 |
| **7994** | 7994 | 12385.622378 | 5192.115385 | 8.013010e-05 | 1.156050e-04 | 9.545691e-05 | 7.889515e-05 | 9.635860e-05 | 1.918419e-04 | 1.000659e-04 | ... | 6.592271e-02 | 4.118145e-02 |
| **7995** | 7995 | 12390.177215 | 5153.939522 | 2.163053e-04 | 2.006127e-04 | 1.394288e-04 | 2.282812e-04 | 1.950765e-04 | 5.304943e-05 | 1.862801e-04 | ... | 7.606965e-02 | 6.082474e-03 |
| **7996** | 7996 | 12389.608696 | 5655.294314 | 2.833072e-02 | 4.516136e-02 | 3.524920e-02 | 3.714351e-02 | 2.777494e-02 | 7.616564e-02 | 4.222060e-02 | ... | 5.101196e-03 | 6.113885e-03 |

7997 rows × 80 columns

#### adata.var

Adata.var returns a dataframe with the gene-related metadata:

| | variable_gene |
|---|---|
| **ATP11A** | True |
| **BCL2** | False |
| **BMP5** | False |
| **CCBE1** | False |
| **CCL21** | False |
| **...** | ... |
| **WNT2** | False |
| **WNT2B** | False |
| **WNT5A** | False |
| **WNT7B** | False |
| **WT1** | False |

140 rows × 1 columns

#### adata.X

Adata.X returns a numpy array with the expression of all genes in all cells, without headers:

```
array([[0., 0., 1., ..., 0., 0., 2.],
       [1., 0., 0., ..., 0., 0., 5.],
       [2., 0., 0., ..., 0., 0., 0.],
       ...,
       [0., 1., 1., ..., 0., 0., 0.],
       [0., 0., 0., ..., 0., 0., 0.],
       [0., 0., 0., ..., 0., 0., 0.]], dtype=float32)
```

##### Filtering and normalizing your cells

The first step of the analysis is to filter out cells that do not show useful expression data. These are typically either cells with too few reads (so they can't be reliably classified), or cells with too many reads that are products of segmentation errors (more than one cell has been captured under the same segmentation region, i.e. doublets).

We can calculate the total number of reads for each cell and save it in .obs as it's done here

```
adata.obs['total_counts']=list(np.sum(adata.X,axis=1))
```

We can then set our filtering threshold. We choose to keep cells only if they have a minimum number of reads (usually anything above 5-7 reads works, but it depends on the quality of the data) and also we should exclude cells with a suspiciously high number of reads, usually >100. These are usually imaging or decoding artifacts, and not real cells.

We now save the raw data in adata.raw and apply normalization and log-transformation to minimize the effect of outliers.The normalization step aims to normalize the number of reads across different cells. There's an ongoing debate in the lab about whether one should normalize or not when using a targeted method like ISS. We believe it sometimes can add artifacts: detection levels for different genes will depend on the intrinsic expression levels, but also variables such as probe efficiencies, or number of probes used per each gene. We suggest trying both normalizing and without normalization, and have a look at the output of the clustering to make a decision.
Log-transformation (which we suggest to perform), is applied to **maximize** the differences between cells with some expression and no expression of a gene, and **minimize** the effect of cells with a very high expression of a certain gene, which helps the clustering.

```
sc.pp.normalize_total(adata, target_sum=1e4)
sc.pp.log1p(adata)
```

We **could** scale the data (scaling is effectively a normalization *by gene*) to give the same importance to all genes in clustering. This is normally done in scRNAseq, but it's not recommended for ISS data: by scaling, we equalize the importance of all genes in clustering. This can lead to overestimating the importance of noise, and worsen the clustering quality: if you have many noisy genes, they will have a higher impact if your data is scaled and, therefore, you will not be able to pick up biologically significant clusters.

#### Principal component analysis, KNN and dimensional reduction.

Next, we apply principal component analysis (PCA) to find the appropriate number of components for dimensionality reduction and clustering. The idea of PCA is to recapitulate the diversity of your data using a reduced set of variables that explain the variability of your original data well enough. This is, on essence, discriminating meaningful variation from residual variation. You can find a deeper explanation in here:
https://builtin.com/data-science/step-step-explanation-principal-component-analysis

After inspecting the PCA result, we can select the top X first principal components to perform a nearest neighbor analysis. We are going to represent our data using graph-based methods, like UMAP. For this, we first need to create a graph by linking each cell to the n closest neighbors (n_neighbors) in the dataset. We can select the numbers of principal components to use. Select “0” to use all.

We can, then, run clustering, by calling either leiden or louvain clustering. By changing the parameter "resolution" we will generate more or less clusters. Higher resolution means more clusters. The clusters (groups of cells) are independent from its visualization, which we will create using UMAP later on.

To VISUALIZE the diversity of our cells, We perform UMAP. The closer two spots are in UMAP, the more similar they will be. With the comand `sc.tl.umap(ad).` we CALCULATE the UMAP embedding. With `sc.pl.umap(umap)` will VISUALIZE the UMAP.

One interesting question we usually have at this stage is "which are the marker genes for the different cell clusters?". To answer this question, we can calculate the differentially expressed genes (DE genes) using the function rank_genes_groups. We first calculate them, indicating our 'key'. Several tests can be run with different properties. For our case, we prefer to use Wilcoxon, but you can use other ones in:https://scanpy.readthedocs.io/en/latest/generated/scanpy.tl.rank_genes_groups.html#scanpy.tl.rank_genes_groups

In the plots, we see that different genes are standing out when comparing specific clusters VS all the cells. Each plot will represent a unique cell cluster, with the genes ranked on the X axis according to their score (Y axis). The higher the score is (Y axis) the more the gene will be differentially expressed in that cluster.

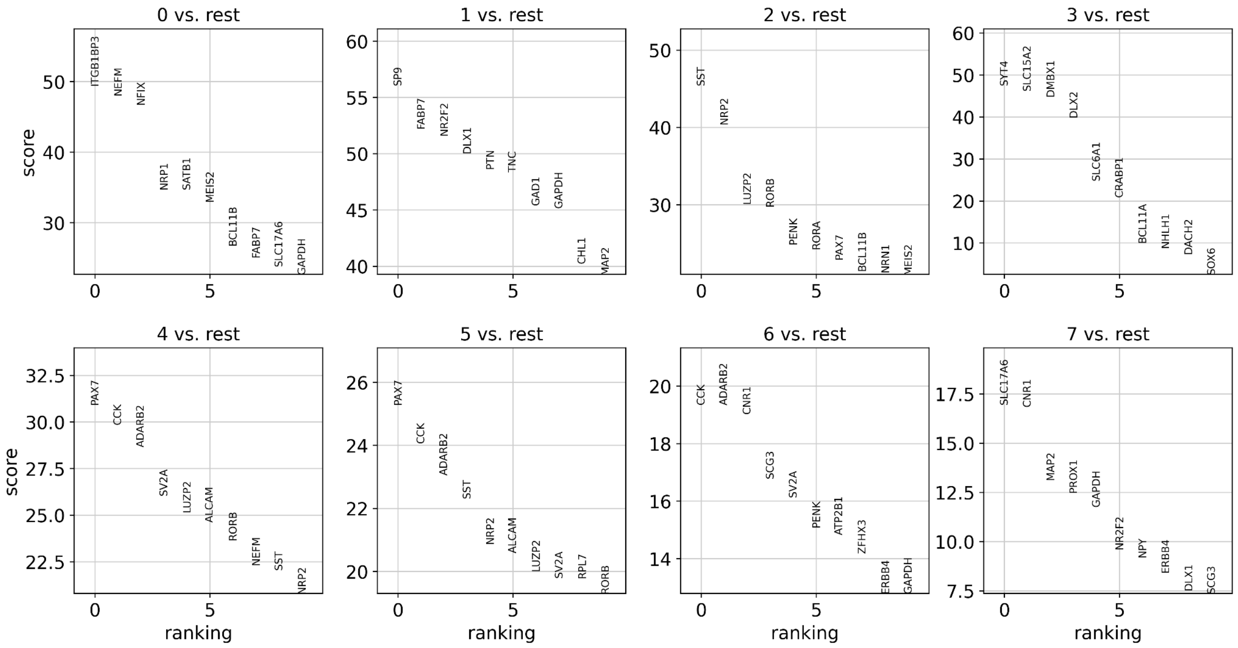


Depending on your data, it might take a while to go through all this analysis successfully. Therefore, it's interesting that you can save and load your processed data whenever you want. To save your object, you use the following command:

```
adata.write(<path>)
```

### Additional functionality in ISS_postprocessing

Once you have generated your annotated data objects, there are multiple things that we can do to interact and visualize the data.

#### Basic plotting function using `sc.pl.spatial`

The first thing that we can do is to plot all of the spatial cells on the basis of their respective x and y coordinates. For this to happen, we need to have the spatial information stored in `adsp.obsm['spatial']` as a numpy array. The way we go about plotting this is to call:

```
sc.pl.spatial(adsp,color='leiden' ,spot_size=100)
```

This will output an image showing you the location of each single cells in space. This will be colored by the column that you specify.

### Advanced features:

### Image deconvolution / denoising (requires GPU)

Our internal tests suggest that it is often beneficial to apply denoising/deconvolution methods to sharpen the images of the rolling circle products, before proceeding to decoding. This is especially advantageous in case of low magnification imaging and/or in signal-crowded tissues, because deconvolution helps separating nearby dots and their respective signals. In case of dense data, it is common to decode 2x or 3x the number of spots compared to non-deconvolved data, and to experience a sharp increase in the decoding quality metrics. To facilitate this, we provide 2 alternative methods for image denoising. Both methods require the use of CUDA-compatible GPUs.

#### Tensorflow-based image deconvolution using Flowdec.

The first method uses the **flowdec** library (https://github.com/hammerlab/flowdec) for tensorflow-accelerated image deconvolution. The library has decent quality documentation, and we suggest you refer to it for advanced tweaking and options. Nevertheless, the library is not currently maintained, and some people have experienced difficulties with it, especially related to how the tiff metadata is handled. We introduced some code to rebuild the tiff metadata, which seems to have fixed the bug, at least for us. In case of difficulties or weird errors, please get in touch with us and we'll do our best to help. Our plan is, in the long run, to move away from flowdec and use something else, but we haven't made up our mind about this yet.

##### Installation and requirements:

Please follow the instructions in the paragraph "Installation of the software toolboxes" for installation guidance.For more advanced installation options and usage details please follow the instructions here:

https://pypi.org/project/flowdec/

**Data formatting and interfacing with the main pipeline**
We have implemented functions that cover the first step of the preprocessing (stacking + maximum projection, intercalated by an image deconvolution step. **If you want to deconvolve your image, you will have to perform the first step of preprocessing in the deconvolution module, and then pipe the projected images into the respective OME_tiff functions depending on the microscope you used.**

*To run image deconvolution using this notebook, you need a CUDA-compatible GPU with functional NVIDIA drivers, cuDNN, etc... and a functional Tensoflow installation.*

**Point spread function**

To do image deconvolution, you will first need to know some parameters about your microscope. Take some time to carefully read the guide below and to collect some information before you attempt image deconvolution. This will save you a lot of computing time and frustration: using the wrong parameters will result in imaging artifacts, so be careful!

These parameters will be used to create a synthetic point-spread function (PSF). If you don't know what a PSF is and why it is important, please read here: https://en.wikipedia.org/wiki/Point*spread*function

You will need to know:

`na` = numerical aperture of the used lens

`m` = lens magnification

`ni0` = refraction index of the immersion medium

`res_lateral` = x,y resolution of the images

`res_axial`: z-resolution of the stack. This is either the spacing between zplanes or the actual z resolution of your lens, whichever is larger. Remember that the actual Z resolution of your lens will match your experiment's resolution only if you acquired the stacks at Nyquist conditions. (https://imb.uq.edu.au/research/facilities/microscopy/training-manuals/microscopy-online-resources/image-capture/nyquist-conditions)

The notebook will guide you through the formatting of data for the creation of your synthetic PSF.

### Image deconvolution of Leica-exported tiffs

##### `deconvolve_leica`

This function organises the raw images from a Leica experiment according to Cycle, Region, Tile, Channel, arranges them in individual 3D stacks, deconvolves each stack and save its maximum projection (default).

The function mirrors the workflow of the `leica_mipping`function from the `ISS_preprocessing` module in the sense that organise the files in a similar way but performs a deconvolution before the maximum projection. The output files and folder have the same name and structure than conventionally projected files. This steps effectively substitutes the `leica_mipping` in preprocessing workflows where image deconvolution is needed.

The function takes the following arguments:

`input_dirs`: This will be a list of the complete paths to the folders containing your imaging cycles. The folders need to be specified in the right order (ie. the first element of the list will be the folder where the first cycle of imaging is saved, and so on) The elements need to be separated by commas. The format for this variable is a `lst` of `str`.

`output_location`: This will be the path where you want the preprocessing output to be saved. Ideally, this should be associated with some type of unique project identifier. The format of this variable is `str`. In case multiple regions are being processed, the funcion will behave differently depending whether a trailing slash `/` is included or not in `output_location`: - if a trailing slash is added, then subfolders for each one of the scanned regions will be created as `_R1`, `_R2`, etc... - if a trailing slash is omitted, then `output_location` will be updated at each region iteration, and each one of the scanned regions will end up in a different `output_location` named as `output_location_R1`,`output_location_R2`, etc...

`image_dimensions`: this is a list of 2 elements [x,y], specifying the x and y sizes of each image (default=[2048, 2048])

`PSF_metadata`: refer to the example above. Metadata for the construction of a synthetic PSF

`chunk_size`: Depending on the GPU you have, you might be or not able to load the entire image stack onto the GPU RAM. If `chunk_size` is unspecified, the function will try to load the entire stack. If your kernel crashes before performing any actual work, the most likely cause is that you are running out of memory. The solution to this RAM shortage is to chunk the stack in sub-stacks, deconvolve each independently and merge them afterwards. A typical `chunk_size` value that works for most GPU is [512,512] in XY. Z is automatically extracted from the data. If your GPU is very small you can try with [256,256] or even [128,128].

`mip`: Specifies if the deconvolved images need to be maximum projected (default = True). If `False` the deconvolved stacks are saved, however we do our ISS analysis in 2D so there's almost never a good reason to save the stack.

`deconvolve_leica` is able to handle multiple regions in the input files, and project them accordingly.

```
from ISS_deconvolution import deconvolution

deconvolve_leica(input_dirs = ['/path/to/cycle1/',
'/path/to/cycle2/',
'/path/to/cycle3/'],
output_location = '/path/to/output_folder/',
PSF_metadata=example_PSF, chunk_size=None, mip=True)
```

### Image deconvolution of CZI files

##### `deconvolve_czi`

This function reads the CZI images from the Zeiss format, gets the images, arranges them in individual 3D stacks, deconvolves each stack and save its maximum projection (default).

The function mirrors the workflow of the `process_czi` function from the `ISS_preprocessing` module in the sense that organise the files in a similar way but performs a deconvolution before the maximum projection. The output files and folder have the same name and structure than conventionally projected files. This steps effectively substitutes the `process_czi` in preprocessing workflows where image deconvolution is needed.

The function takes the following arguments:

`input_file`: the path to the CZI file that you want to preprocess, down to the czi file (included)

`outpath`: the folder where you want to save the maximum-projected images. Ideally this would be a `/mainoutputfolder/region/preprocessing/mipped/`folder structure, for consistency with our way of organising the data.

`cycle`: here you have to manually specify to which ISS cycle the images refer to. This is a `int` number, where 1 refers to cycle 1 and so on. If `cycle=0` the function will not work.

`tile_size_x` and `tile_size_y`: these refer to the size in pixel of your camera field of view. Most cameras are 2048x2048, so that's the default if you don't specify them, but adjust them if your camera has a different field of view size.

`chunk_size`: Depending on the GPU you have, you might be or not able to load the entire image stack onto the GPU RAM. If `chunk_size` is unspecified, the function will try to load the entire stack. If your kernel crashes before performing any actual work, the most likely cause is that you are running out of memory. The solution to this RAM shortage is to chunk the stack in sub-stacks, deconvolve each independently and merge them afterwards. A typical `chunk_size` value that works for most GPU is [512,512] in XY. Z is automatically extracted from the data. If your GPU is very small you can try with [256,256] or even [128,128].

`mip`: Specifies if the deconvolved images need to be maximum projected (default = True). If `False` the deconvolved stacks are saved, however we do our ISS analysis in 2D so there's almost never a good reason to save the stack.

**Keep in mind that `deconvolve_czi` can only accept 1 file at the time, unlike `deconvolve_leica`. This means that you need to deconvolve ONE cycle at the time, and manually specify the cycle number in the function for appropriate naming of the output files. You can also embed this function in a loop to process multiple cycles if you want, but make sure to pass the right arguments.**

```
from ISS_deconvolution import deconvolution

deconvolve_czi(input_file, outpath, image_dimensions=[2048, 2048],
PSF_metadata=example_PSF, chunk_size=None, mip=True, cycle=0, tile_size_x=2048,
tile_size_y=2048)
```

#### ML-accelerated image restoration using 2D CARE:

##### Background

Content Aware Image Restoration (CARE) is a ML-based method for image denoising/deconvolution/restoration, please have a look at:
https://csbdeep.bioimagecomputing.com/
When appropriately trained, it can be used as a very fast alternative to image deconvolution. Its main advantage is that <u>it can work directly on projected images</u>, significantly reducing the computing requirements. This speeds up the denoising process by a factor of at least a 100 compared to Flowdec (which is already a faster method than most available), although we acknowledge it is likely less accurate than "true" deconvolution. From our benchmarking, CARE seems to be the way to go for most experiments. We advise the user to do a test run on a small

sample with both wlowdec and CARE, compare the results and proceed using CARE only if the results are satisfyingly similar.

##### Formatting and interfacing with the main pipeline

2D CARE should be applied after the preprocessing step. **The input files needed for CARE are those in the *preprocessing/ReslicedTiles/* folder (projected resliced tiles arranged by channel)**.

By default our notebook excludes the DAPI images from the CARE-based deconvolution process when using our pre-trained models: these models have not been trained on DAPI and might produce weird artifacts in the nuclei shapes. For this, the user needs to specify the channel number corresponding to the DAPI signal, so the corresponding images are excluded from the CARE workflow. The unprocessed DAPI images and the metadata are automatically copied into the destination folder.

##### Apply a pre-trained model

For convenience, we provide a pre-trained model which works fairly well in our hands. The model has been trained using a set of images in all channels (except DAPI) coming both from Leica and Zeiss, using data acquired using 20x and 40x objectives, and spanning a range of tissues and model organisms.

The script to run to apply the model is:

```
ISS_CARE(directory = '/path/to/experiment/preprocessing/ReslicedTiles/',
             basedir_model = '/path/to/models/',
             model_name = 'spots',
             output_dir = '/path/to/output_folder',
             DAPI_ch=5)
```

##### Explanation of the variables

`directory` = type: `str`. this is the path to your `/ReslicedTiles/` folder containing the raw mipped images to deconvolve.

`basedir_model` = type: `str`. The path to the folder containing the pre-trained models.
`model_name` = type: `str`. The name of the model to apply.

`output_dir` = type: `str`. This is the output directory for the denoised images. A `/preprocessing/ReslicedTiles/` folder will be created here, along with specific subfolders reflecting the folder tree of the input path.

`DAPI_ch` = type: `int`. Number of the channel containing the DAPI images. This is to exclude DAPI from the CARE processing.

This notebook will output, in the chosen output folder, the transformed images, ready for the downstream analysis. It will also copy, to the output folder, the DAPI unprocessed images as well as the tilepos.csv file, containing the xy coordinates of the tiles. These files should be used as input for the SpaceTx formatting function, and move on from there.

##### Train your own model

The csbdeep package (which is the core of CARE) has excellent documentation about how to train and evaluate models for an image denoising task. We provide scripts to train your own models and to package your data for training. These scripts are just the demo scripts coming with the csbdeep package. Please refer to the following links for a detailed guide.

If you want to train your own model, or add training to an existing one, you should refer to:
https://csbdeep.bioimagecomputing.com/doc/datagen.html
This allows you to package data for training, and it requires you to provide pairs of noisy/deconvolved projected images. They have to have the same name and have to be EXACTLY registered.
Once the data is properly formatted, then you should run the code described here:
https://csbdeep.bioimagecomputing.com/doc/training.html
This actually trains and saves your model for later use. WARNING: this can be very slow, depending on the size of the training data, and the power of your GPU.

And finally, after you trained your model, you can apply it to new images to denoise them using the Jupyter notebook above (WARNING: you should NEVER apply the model on the same images you used for training, this violates the rule number one of machine learning).

### References

Biancalani, Tommaso, Gabriele Scalia, Lorenzo Buffoni, Raghav Avasthi, Ziqing Lu, Aman Sanger, Neriman Tokcan, et al. 2021. “Deep Learning and Alignment of Spatially Resolved

Single-Cell Transcriptomes with Tangram.” *Nature Methods* 18 (11): 1352–62.
Czech, Eric, Bulent Arman Aksoy, Pinar Aksoy, and Jeff Hammerbacher. 2019. “Cytokit: A Single-Cell Analysis Toolkit for High Dimensional Fluorescent Microscopy Imaging.” *BMC Bioinformatics* 20 (1): 448.
Gyllborg, Daniel, Christoffer Mattsson Langseth, Xiaoyan Qian, Eunkyoung Choi, Sergio Marco Salas, Markus M. Hilscher, Ed S. Lein, and Mats Nilsson. 2020. “Hybridization-Based in Situ Sequencing (HybISS) for Spatially Resolved Transcriptomics in Human and Mouse Brain Tissue.” *Nucleic Acids Research* 48 (19): e112.
Ke, Rongqin, Marco Mignardi, Alexandra Pacureanu, Jessica Svedlund, Johan Botling, Carolina Wählby, and Mats Nilsson. 2013. “In Situ Sequencing for RNA Analysis in Preserved Tissue and Cells.” *Nature Methods* 10 (9): 857–60.
Muhlich, Jeremy L., Yu-An Chen, Clarence Yapp, Douglas Russell, Sandro Santagata, and Peter K. Sorger. 2022. “Stitching and Registering Highly Multiplexed Whole-Slide Images of Tissues and Tumors Using ASHLAR.” *Bioinformatics* 38 (19): 4613–21.
Qian, Xiaoyan, Kenneth D. Harris, Thomas Hauling, Dimitris Nicoloutsopoulos, Ana B. Muñoz-Manchado, Nathan Skene, Jens Hjerling-Leffler, and Mats Nilsson. 2020. “Probabilistic Cell Typing Enables Fine Mapping of Closely Related Cell Types in Situ.” *Nature Methods* 17 (1): 101–6.
Stringer, Carsen, Tim Wang, Michalis Michaelos, and Marius Pachitariu. 2021. “Cellpose: A Generalist Algorithm for Cellular Segmentation.” *Nature Methods* 18 (1): 100–106.
Weigert, Martin, Uwe Schmidt, Tobias Boothe, Andreas Müller, Alexandr Dibrov, Akanksha Jain, Benjamin Wilhelm, et al. 2018. “Content-Aware Image Restoration: Pushing the Limits of Fluorescence Microscopy.” *Nature Methods* 15 (12): 1090–97.
